# Multimodal spatial alignment and morphology mapping with MOSAICField

**DOI:** 10.64898/2026.03.10.710931

**Authors:** Xinhao Liu, Hongyu Zheng, Peter Halmos, Julian Gold, Erik Storrs, Li Ding, Benjamin J. Raphael

## Abstract

Recent efforts to build comprehensive tissue and tumor atlases leverage diverse spatial technologies to measure transcriptomic, proteomic, epigenetic, and other modalities with hundreds to thousands of features at thousands to millions of spatially resolved locations in a tissue slice. Integrating such data across spatial technologies that differ in molecular features, spatial resolution, and tissue morphology remains a major challenge. We introduce MultimOdal Spatial Alignment and Integration with Coordinate neural Field (MOSAICField), a unified framework for aligning spatial slices across arbitrary combinations of experimental modalities. MOSAICField computes two types of spatial alignments across multiple slices from the same tissue: *physical alignment*, which reconstructs a contiguous 3D model of the original tissue, and *morphological alignment*, which maps distinct morphological or anatomical structures, such as ducts, veins, or neurons, that may traverse the tissue at different angles relative to the direction of slicing. MOSAICField computes both alignments using a deep neural network that estimates a nonlinear deformation field with a multimodal feature loss. We evaluate MOSAICField on simulated data and a prostate cancer sample from the Human Tumor Atlas Network (HTAN), containing more than a dozen spatial slices with multimodal profiling data. MOSAICField constructs an accurate 3D tumor model, tracks the architecture of the prostatic ductal system, and improves analysis of features within and across modalities, outperforming existing methods.

## 1 Introduction

Spatial sequencing technologies such as spatial transcriptomics [44, 53], spatial proteomics [38], and spatially resolved epigenomics [15, 37] provide high-throughput measurements of mRNA and protein expression, epigenetic changes, and other modalities, at cellular and subcellular resolution. While there are recent successes in simultaneous spatial profiling of multiple modalities on the same tissue [35, 66], most of the widely available spatial technologies measure only one single modality. Atlas projects such as the Human Tumor Atlas Network (HTAN) apply different technologies to different slices from the same tissue, extending cell atlas consortiums such as Human Cell Atlas (HCA) to both spatial and multimodal measurements.

Integrating data from multiple spatial slices enables *in silico* reconstruction of 3D tissue models, opening new ways to understand three-dimensional tissue microenvironments and their spatial dynamics [42]. Integrating spatial data from different modalities provides a more comprehensive characterization of the spatial heterogeneity of cell types and cell states within a tissue [33]. A number of tools exist for single-cell multi-omics integration [2, 3, 8, 14, 30, 68], but ignore spatial coordinates for linking measurements across modalities. On the other hand, multiple methods have been developed to register spatial slices of the same modality [11, 25, 29, 34, 65, 69]. However, these methods are not easily extensible to cross-modality alignment. Some methods [10, 13, 70] integrate spatial multiomics into a common embedding space, ignoring spatial structure of the data. Recently, two methods, CAST [56] and SANTO [32], were introduced for spatial multi-omics alignment. CAST employs graph neural networks and the B-splines [51], while SANTO uses dynamic graph convolutional neural networks [61] to learn similar omics features across slices. However, both methods rely on the expression of common genes across slices; the multi-omics aspect of the two methods largely refers to cross-platform integration of spatial transcriptomics data, such as aligning STARmap [60] to MERFISH [67] or 10x Genomics Visium to 10x Genomics Xenium. The utility of these methods on cross-modality alignment – e.g. spatial transcriptomics and H&E images – has not been demonstrated.

We introduce MOSAICField (**M**ultim**O**dal **S**patial **A**lignment and **I**ntegration with **C**oordinate Neural **Field**), a method to align, register, and track morphology for spatial slices across multiple modalities and resolutions (Fig. 1). Unlike existing methods, MOSAICField aligns multimodal spatial slices without common features. Given a stack of slices from the same tissue, each potentially from a different modality, MOSAICField aligns each pair of adjacent slices and composes these alignments for a multimodal 3D view of the tissue.

**Fig. 1:**
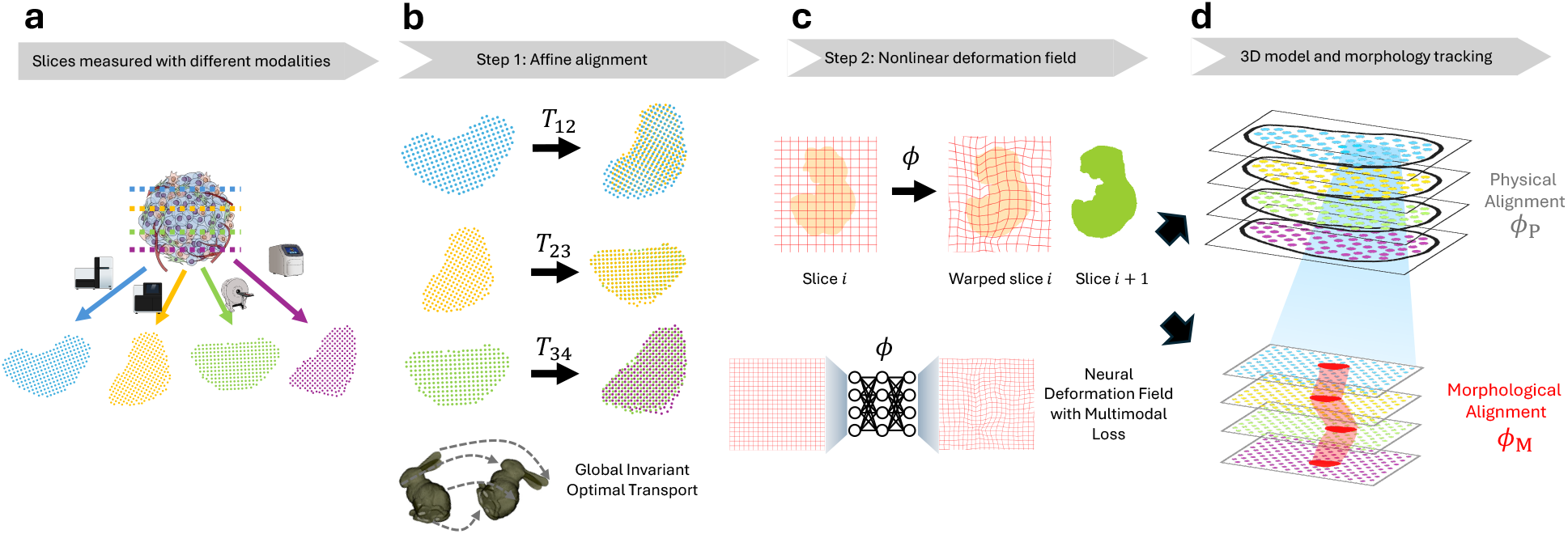
Overview of MOSAICField. **a**, Multiple slices from a 3D tissue are assayed using different spatial profiling or imaging technologies, e.g. spatial transcriptomics, spatial proteomics, H&E staining, etc. **b**, In step 1, MOSAICField uses global invariant optimal transport to compute an affine transformation **T** between each pair of adjacent slices regardless of their modalities. The affine transformations are used as initializations for nonlinear alignment. **c**, In step 2, MOSAICField learns a nonlinear deformation field between each pair of affine-aligned slices with deep neural networks to capture nonlinear distortions. **d**, Learning a nonlinear transformation at global scale, MOSAICField computes a physical alignment that reconstructs a multimodal 3D model of the tissue. Learning a nonlinear transformation at local scale, MOSAICField computes a morphological alignment that aligns morphological structures (e.g. ducts, veins) that traverse the 3D tissue in arbitrary directions, including directions not perpendicular to the direction of slicing.

MOSAICField distinguishes between two distinct goals of spatial alignment: (1) *physical alignment*, or the registration of slices into a common coordinate system describing the original 3D organization of the slices; (2) *morphological alignment*, or the identification and tracking of specific morphological features – such as ducts, vascular networks, or tumor boundaries across the physically aligned slices in non-perpendicular directions to the direction of slicing (Fig. 1d). Physical alignment restores the spatial context in the original tissue and corrects for spatial distortion, scaling, rotation, and translation across 2D slices to ensure each slice is accurately placed relative to others, allowing the entire dataset to form a coherent 3D representation of the original tissue. Morphological alignment tracks and analyzes morphological structures within the reconstructed 3D model. For example, the human prostate gland comprises a network of ductal system that serves as conduits for prostatic secretions [40], and morphological alignment is needed to track this complicated architecture within a 3D tissue. Similarly, tracking morphology is essential to study ductal cancers such as ductal carcinoma in situ (DCIS) or invasive ductal carcinoma (IDC). Most existing alignment methods do not distinguish between physical and morphological alignment, either assuming the two are identical, or using manual annotations to perform morphological tracking [26].

MOSAICField starts by computing an affine alignment between two multimodal slices using an extension of global invariant optimal transport (OT) [1, 16] (Fig. 1b). From the affine aligned coordinates, MOSAICField computes both physical and morphological alignments as nonlinear mapping based on the concept of neural implicit representations of deformation fields [6, 55, 58] and multimodality image registration [20, 49] (Fig. 1c).

We validate MOSAICField on a simulated dataset and a multimodal prostate cancer sample from HTAN. MOSAICField corrects for global distortions and achieves reliable morphology tracking in simulation, outperforming previous methods. On the multimodal prostate cancer dataset, MOSAICField integrates slices from three different spatial platforms (10x Genomics Xenium, H&E histological staining, multiplex imaging using Co-Detection by Indexing (CODEX)) into one uniform multimodal 3D model of the sample and accurately tracks the path and shape of the prostate ductal system in 3D. Finally, we show that MOSAICField infers functional relationships between multimodal features, such as gene and protein expression, enabling the study of complex biological interactions using multimodal information within the tissue microenvironment.

## 2 Methods

Given spatial profiling data from multiple slices from the same tissue – potentially using multiple different profiling modalities – MOSAICField aligns each pair of adjacent slices. Let 𝒟 **=** {(**X**^(1)^, **S**^(1)^, …, **X**^(*M*)^, **S**^(*M*)^)} be spatial datasets from *M* tissue slices, where **X**^(*i*)^ is an *n*_*i*_ *m*_*i*_-dimensional feature matrix for *m*_*i*_-dimensional features of *n*_*i*_ locations on the slice *i*, and **S**^(*i*)^ is the *n*_*i*_ 2-dimensional spatial coordinate matrix. Both the definition of a location and the dimension of a location’s feature depend on the specific modality. For example, for single-cell spatial transcriptomics technologies a location corresponds to a cell and a feature is a gene expression vector for a cell; for spatial imaging data, a location is a pixel and a feature is the channel intensities of that pixel. For a pair of slices possibly from different modalities, MOSAICField aims to find a nonlinear transformation registering one slice onto the other. MOSAICField first computes an affine transformation **T** : ℝ^2^ → ℝ^2^, then, from the affine-aligned coordinates of the two slices, learns a nonlinear deformation *ϕ* : ℝ^2^ → ℝ^2^ via neural fields.

When aligning serial sections from the same tissue, there are two distinct objectives. First, one aims to reconstruct the original 3D architecture of the specimen by positioning all slices *in-silico* according to their physical positions in the original tissue. Second, one seeks to trace continuous morphological or anatomical structures across sections within this reconstructed 3D model. These objectives require distinct alignment formulations. The first emphasizes spatial consistency along the slicing (*z*) axis, aligning locations that were physically adjacent in the original tissue. The second emphasizes the continuity of structures that may intersect the slicing plane at arbitrary orientations. For example, a duct traversing the organ obliquely should be aligned across sections according to its morphological trajectory, not its *z*-axis proximity. We define two types of alignment problems to address these objectives.

### Definition 1 (Physical alignment)

*A physical alignment ϕ*_*P*_ *registers all spatial slices into a common coordinate system that recapitulates their original 3D spatial organization*.

### Definition 2 (Morphological alignment)

*A morphological alignment ϕ*_*M*_ *aligns corresponding morphological structures that may traverse the tissue at non-perpendicular orientations relative to the slicing direction*.

A major challenge is to distinguish these two alignments given only the slices themselves. One straight-forward distinction from a mathematical perspective is to constrain the physical alignment to an affine transformation while allowing nonlinear transformations for morphological alignment, as morphological structures often involve complex local deformations. However, affine functions are typically insufficient for reconstructing real 3D tissues, where sample preparation can introduce nonlinear distortions due to uneven shrinkage, tearing, non-parallel slicing, or inconsistent sectioning. Therefore, we define both alignments to be nonlinear, denoting physical alignment as *ϕ*_*P*_ and morphological alignment as *ϕ*_*M*_. Importantly, once nonlinearities are allowed for both alignment types, the distinction between them becomes less clear mathematically. Any threshold distinguishing “small” versus “large” nonlinearities is inherently arbitrary. The solution in MOSAICField is to distinguish physical vs. morphological alignment based on allowing a fixed “amount” of nonlinearity (by regularizing *ϕ*) over different spatial lengths. Specifically, we first compute an affine transformation **T** (§2.1) and then compute separate *ϕ*_*P*_ and *ϕ*_*M*_ using neural fields (§2.2) over different physical scales (§2.3).

### 2.1 Step 1: Affine alignment

#### Objective

Given slices 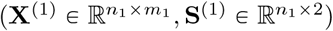 of modality 1 and 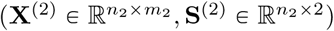 of modality 2, MOSAICField computes an affine transformation **T** : ℝ^2^ → ℝ^2^ that minimizes the following objective function:

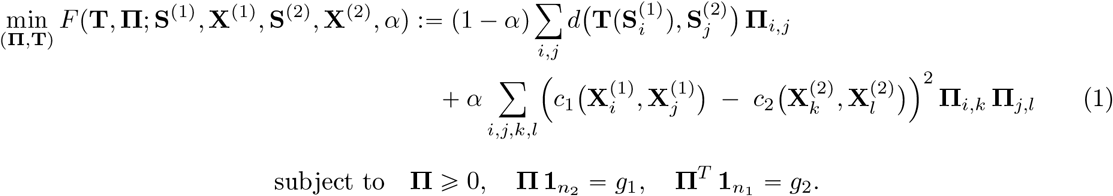

Here, 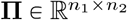 is a coupling matrix between the locations of the two slices. *α* is a hyperparameter controlling the contribution of the two loss terms with *α* P r0, 1s, *g*_1_ and *g*_2_ are uniform probability distributions over the locations in each slice, *d*(*x, y*) is the Euclidean distance between two 2-dimensional coordinates *x* and y. 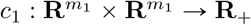 is a feature distance function that measures the distance between two features in the *m*_1_-dimensional space for the features of slice 1, and similarly for *c*_2_.

The objective function *F* is a sum of two terms: the first term (Wasserstein term) evaluates the distance between locations in two slices transformed by **T**, while the second term (Gromov-Wasserstein term) evaluates the difference in features across pairs of aligned locations in each slice. This objective function is an extension of Fused Gromov-Wasserstein optimal transport (OT) [57], which was introduced for spatial alignment in PASTE [34, 65] and used in later OT-based alignment methods [17, 29], but has two important differences. First, the role of these terms is opposite to that of existing OT-based alignment methods, with the Gromov-Wasserstein term [47] allowing comparison of features across different metric spaces and thus enabling alignment of slices from completely different spatial modalities (e.g. RNA expression, protein abundance, H&E staining). Second, we model global geometric invariances by including the affine transformation **T** in the Wasserstein term for spatial coordinates. Similar formulations have been explored in the context of representation learning [16] and computer vision [12, 48] as a global invariant optimal transport problem [1, 16].

MOSAICField solves for **Π** and **T** simultaneously using a block coordinate descent algorithm (Supplement §S1) that alternates between learning a coupling **Π** by solving Fused Gromov-Wasserstein Optimal Transport, and learning the parameter **T** by solving a least squares optimization problem similar to generalized Procrustes analysis [59].

### 2.2 Step 2: Nonlinear alignment via neural field

#### Problem Formulation

The goal of this step is to compute a nonlinear transformation *ϕ* that describes nonlinear deformations between slices. We assume the two slices are already affine **T**-aligned in this step so that *ϕ* learns residual nonlinearities of **T**. In contrast to step 1, where each slice is represented as a point cloud **X, S**, here we treat each slice as a multi-channel image to leverage the extensive literature on deformable image registration [4, 5, 43, 52, 55, 63]. This formulation is well motivated, as most experimental modalities inherently produce image-based data [26, 62] or produce readouts on a grid-like structure [45], and many non-imaging modalities can be readily rasterized into image representations [11].

Let 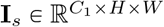 and 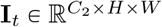 be the affine-aligned source and target images, with *C*_1_ and *C*_2_ channels, respectively. The objective is to find a nonlinear function *ϕ* : ℝ^2^ → ℝ^2^ that aligns **I**_*s*_ onto **I**_*t*_ in a “minimal” way. We solve the unsupervised image registration problem by minimizing the following loss function:

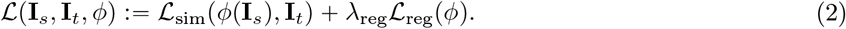

Here, ℒ_sim_ is a similarity loss that measures the distance between target **I**_*t*_ and the warped source image ℒ (**I**_*s*_), and ℒ_reg_ is a regularization term on *ϕ* with weight *λ*_reg_. In this work, we learn *ϕ* by learning a displacement function **v**_*θ*_ : ℝ^2^ → ℝ^2^ and let *ϕ* (*x*) *= x +* **v**_*θ*_ (*x*). We use a multi-layer perceptron (MLP) neural network to model **v**_*θ*_, which takes a 2D coordinate *p* as input and outputs the 2D displacement vector at that position. We use *θ* to denote the neural network parameters. The following sections describe construction of ℒ_sim_ and ℒ_reg_.

#### Generalized Modality Independent Neighborhood Descriptor

Measuring similarity of images in a multimodal setting, with different number of channels and channels representing different aspects of data is very challenging. For example, for spatial transcriptomics, each channel typically corresponds to spatial expression of a gene, while for histology images the channels represent the intensity of different colors under bright field microscopy. To design a suitable ℒ_sim_ for our task, we draw inspiration from the seminal work of Modality Independent Neighborhood Descriptor (MIND) [20, 21] proposing a neighborhood descriptor for single-channel multimodal images. We introduce a version of MIND that generalizes to multimodal images with any number of channels. Specifically, let *ρ* be a patch radius such that each image patch is defined as 𝒫 = {−*ρ*, −*ρ* +1,, *ρ* − 1, *ρ*} × {−*ρ*, −*ρ* +1,, *ρ* ℒ1, *ρ*} with size (2*ρ* +1)^2^. Additionally, let 𝒩 ⊂ ℤ^2^ be a neighborhood around origin with size *k*, e.g. 𝒩 *=* {(ℒ 1, 0), (0, ℒ 1), (1, 0), (0, 1)} with *k*” 4. For each pixel *p* in each image **I**, we define its *patch descriptor D*′^**I**^(*p*) ∈ ℝ^*k*^ as a vector of length *k*, indexed by elements in 𝒩 :

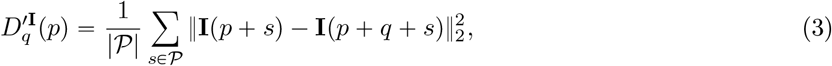

for each neighboring pixel *p + q* with *q* ∈ 𝒩. Here **I** *p* means the channel values at pixel *p*. Then, we normalize the patch descriptor of each pixel to arrive at the generalized modality independent neighborhood descriptor *D*^**I**^(*p*) ∈ ℝ^*k*^ for that pixel:

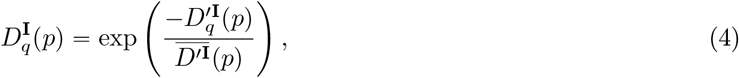

where 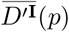 is the average over the *k*-long patch descriptor: 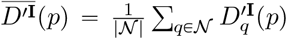. Finally, for the warped source and target images *ϕ* (**I**_*s*_), **I**_*t*_, we define their neighborhood similarity loss by computing the difference over all neighborhood descriptors of all pixels across the two images:

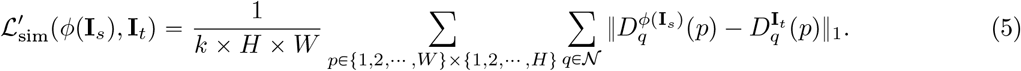

Intuitively, the modality independent neighborhood descriptor *D*^**I**^ *p* encodes the neighborhood structure surrounding pixel *p* within one modality, and its elements describe the difference between the patch around *p* and the patch around *p + q*, for each offset *q*. The neighborhood similarity loss 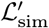 compares the local patch descriptors, where two pixels on different modality are considered similar if the differentials of these pixels against their neighbor show a similar spatial pattern. This enables direct comparison of local similarity without the need of cross-modality measurements. On the other hand, our proposal is connected to a locality restricted version of Gromov-Wasserstein optimal transport problem, from which we can arrive at similar formulations. We discuss these connections in Supplement §S3.

In practice, we add a second, single-channel component. We first reduce each image to a single channel by averaging all channel intensities, scale the values via z-score normalization to account for variations across modalities, then calculate Mean Squared Error (MSE) loss over the normalized average intensity of warped source image *ϕ*(**I**_*s*_) and target image **I**_*t*_. Combining both components:

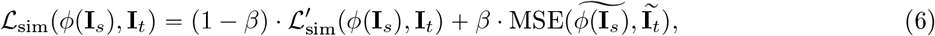

where Ĩ denotes the z-score normalized grayscale version of image **I**, and *β* is a hyperparameter balancing the contribution of the two terms.

#### Regularization and Optimization

We implement local Jacobian regularization as well as magnitude regularization. See Supplement §S2 for details.

### 2.3 Physical and Morphological alignment

As discussed above, distinguishing between “physical nonlinearities” and “morphological nonlinearities” are difficult mathematically. Our solution is to apply the same MOSAICField nonlinear deformation field for both alignments but at two different scales: global and local. Given two slices 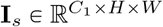 and 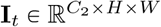 (already affine-aligned as required by our non-linear algorithm), we define the physical alignment *ϕ*_*P*_ as obtained by applying the MOSAICField deformation field on a global scale. Since modern whole-slide imaging platforms often generate gigapixel-resolution images and it is infeasible to run our algorithm on full-resolution data, we apply MOSAICField to low-resolution whole-slide images and interpolate back to full-resolution (Supplement §S5.1). We say that *ϕ*_*P*_ (**I**_*s*_) is physically aligned with **I**_*t*_ this way. The *ϕ*_*P*_ between adjacent slices in a stack can then be composed to reconstruct the overall 3D model of the tissue.

With *ϕ*_*P*_ (**I**_*s*_) and **I**_*t*_ physically aligned, we focus on a local region 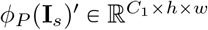 and 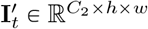 for morphological alignment, where *h < H* and *w < W*. We define the morphological alignment *ϕ*_*M*_ as obtained by applying MOSAICField on this local but full-resolution scale after physical alignment (i.e., we run the nonlinear algorithm in §2.2 on *ϕ*_*P*_-aligned slices). Thus, *ϕ*_*M*_ (*ϕ*_*P*_ (**I**_*s*_)) is morphologically aligned with **I**_*t*_, with *ϕ*_*M*_ learning to map the detailed morphological structures at a local, high-resolution scale not captured by affine and physical alignments.

In this two-scale formulation, MOSAICField models hierarchical aspects of tissue organization: at the global, low-resolution scale, *ϕ*_*P*_ emphasizes overall geometric consistency across slices while paying less attention to local morphology changes; at the local, full-resolution scale, the expressive power of the neural network *ϕ*_*M*_ captures fine-grained nonlinear local distortions and morphological variations.

## 3 Results

### 3.1 Evaluation on simulated spatial multimodal data

We evaluate MOSAICField against four recent spatial alignment methods—STalign [11], PASTE [65], SANTO [32], and CAST [56]—under two simulated multimodal scenarios designed to test physical and morphological alignment, respectively. The first simulation tests robustness under global geometric distortions. We use a Stereo-seq E10.5 mouse embryo slice [9] as the target slice and generate a source slice by applying realistic distortions that mimic experimental artifacts from tissue preparation, mounting, and segmentation. Specifically, we apply a smooth sinusoidal deformation to the spatial coordinates (Supplement §S4), scale the slice by 1.1 ×, and rotate it clockwise by 25 degrees (Fig. 2a). To simulate multimodal features, we partition the original gene set into two disjoint halves: one assigned to the source and the other to the target, ensuring that no genes are shared. We compute PCA embeddings within each slice’s feature space and apply a random linear projection to the source features (Supplement §S4).

**Fig. 2:**
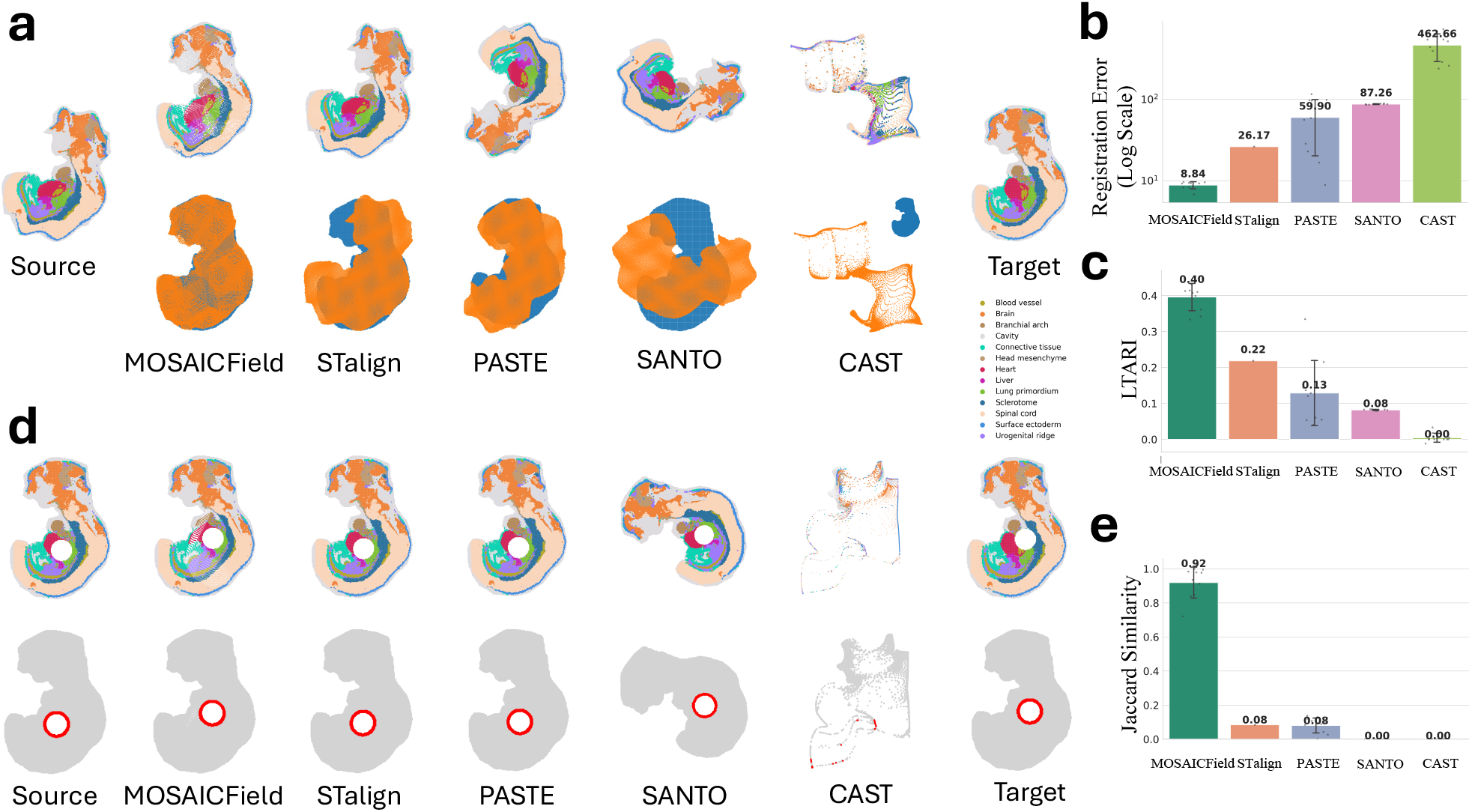
MOSAICField accurately infers physical and morphological alignments on simulated spatial multimodal slices. **a**, Simulated multimodal slices derived from an E10.5 mouse embryo. The source slice is globally distorted from the target to simulate the *physical alignment* problem. Alignment results are shown for MOSAICField, STalign, PASTE, SANTO, and CAST. **b**, Registration error of each method for the physical alignment in **(a)** with error bar. **c**, Label transfer ARI for the physical alignment in **(a)** with error bar. **d**, Simulated multimodal slices illustrating the *morphological alignment* problem. A hollow lumen is embedded at different positions in the source and target slices. The alignment results of all five methods are shown; the red band highlights the ring of cells surrounding the lumen in each slice. **e**, Jaccard similarity between the red bands in each method’s aligned source slice and the target slice with error bar. Error bars indicate standard deviation over 10 independent runs of each method.

We apply all five methods to align the distorted source slice to the target. MOSAICField accurately recovers both the global rotation and scaling, and its nonlinear deformation field effectively corrects the sinusoidal distortions, yielding near-perfect overlap between source and target (Fig. 2a). In contrast, STalign and PASTE fail to recover the correct rotation: STalign produces a small misalignment, while PASTE flips the slice entirely due to its rigid shape-matching formulation. SANTO and CAST, which rely heavily on shared features across slices, fail to produce meaningful alignments under the truly multimodal setting where no direct feature correspondence exists. We quantify the alignment accuracy using two metrics, the registration error and the label transfer adjusted Rand index (LTARI) [34]. The registration error is defined as the average distance between each cell on the aligned source slice and its ground truth corresponding cell on the target slice, and the LTARI measures how well the alignment respects cell type annotations (Supplement §S4). To quantify variability, we perform 10 independent runs for each method and report sample standard deviation as error bars for each evaluation metric. MOSAICField achieves the lowest registration error (Fig. 2b) and the highest LTARI (Fig. 2c), indicating that MOSAICField aligns each source cell to the correct target, and the alignment is biologically reasonable as it matches cell type annotations across slices.

The second simulation evaluates each method’s ability to align morphological structures that traverse tissue slices obliquely, testing MOSAICField ‘s morphological alignment capability. Starting from the same E10.5 embryo slice, we embed a hollow circular structure within the embryo to represent a lumen (Fig. 2d). In the source slice, this lumen is translated such that it partially overlaps with the target’s lumen, simulating a non-perpendicular trajectory. The multimodal features are generated in the same way as the previous step. Notice that while this simulation is not strictly on a local scale, since there is no global distortion and the slices differ only in the lumen’s position, this is a morphological alignment problem. We assess performance using the Jaccard similarity between the rings of cells surrounding the lumen in the aligned source and target slices (Fig. 2e). MOSAICField achieves the highest Jaccard similarity, demonstrating its ability to correctly track and align continuous morphological structures across slices. Competing methods either incompletely match the luminal region or fail entirely under the combined challenge of morphology variation and multimodality.

Additionally, we conduct ablation studies to establish contributions of various components to the performance of MOSAICField (§ S4.4). We find that both the multimodal MIND loss and the single-channel MSE loss are essential, and that MOSAICField is robust to a wide range of parameter combinations (Fig. S2).

We emphasize that the four competing methods tested here – even those that claim to analyze multimodal data – are not designed for the type of multimodal data that we simulate here. STalign and PASTE were designed and tested only on spatial transcriptomics data in their published versions. SANTO and CAST are described as multimodal alignment methods; however, they require a subset of common features across slices, and thus do not generalize to multimodal data from drastically different platforms that do not share features (e.g. transcriptomics and H&E images) which we evaluate here.

### 3.2 MOSAICField physical alignment of prostate cancer samples

We apply MOSAICField to prostate sample HT891Z1 of the multimodal 3D spatial dataset of HTAN [54]. We analyze 16 consecutive slices labeled Z0 to Z215 in a region transitioning from normal glands to Gleason Pattern 3 (GP3) prostate adenocarcinoma, with the number following Z indicating the distance in *µ*m from the top of the tissue. The 16 slices were measured using three different spatial platforms: Z0, Z100, Z150, Z215 with single-cell spatial transcriptomics using the 10x Genomics Xenium platform; Z10, Z20, Z25, Z30, Z135, Z140, Z145, Z170 with histological staining (H&E); and Z65, Z155, Z195, Z210 with multiplex imaging of proteins using Co-Detection by Indexing (CODEX) (Fig. 3a). Thus, 9 out of 15 adjacent pairs have different modalities. The Xenium panel includes 475 genes. The CODEX panel includes 25 proteins. Each slice was prepared separately, and technical differences across experiments caused tissue rotation, translation, and stretching across slices so that the positions of adjacent slices are not directly comparable (Fig. 3b), necessitating physical alignment.

**Fig. 3:**
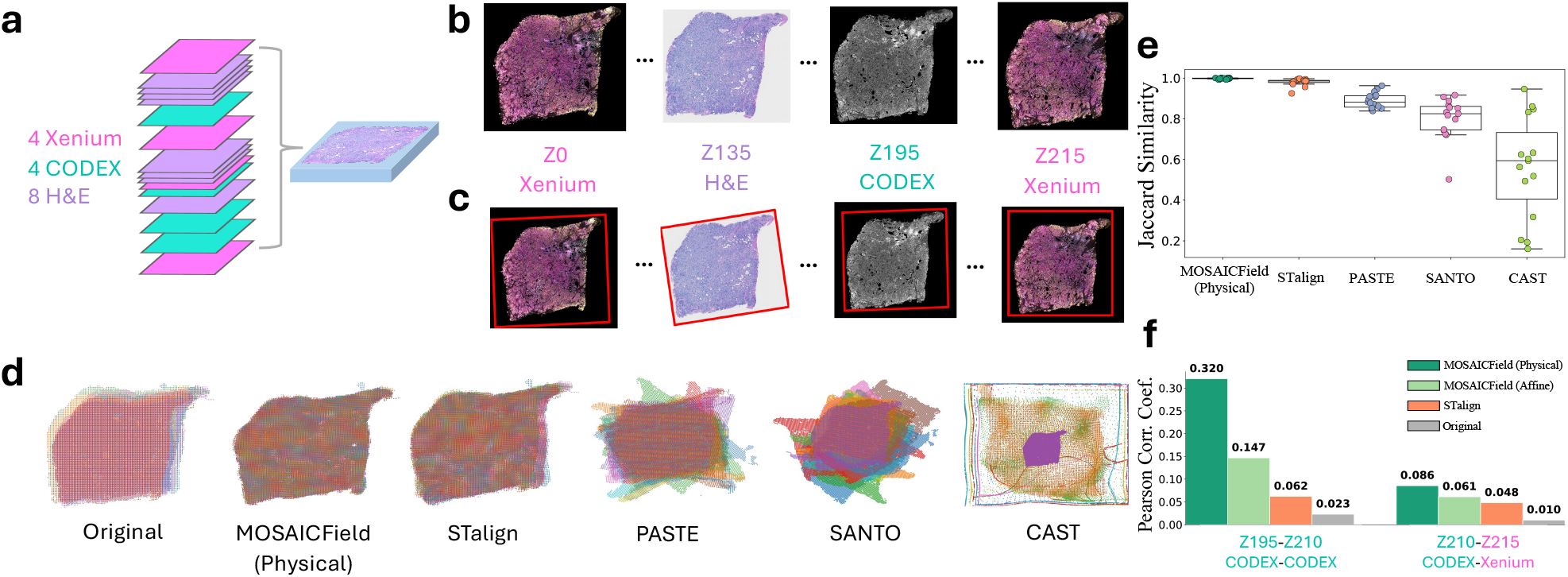
MOSAICField physical alignment of prostate cancer slices. **a**, 16 slices from prostate cancer sample HT891Z1 were assayed using three different experimental technologies: 10x Genomics Xenium transcriptomics, CODEX, and H&E staining. **b**, An example of the raw, unaligned images from four slices. **c**, The images of the slices after MOSAICField affine alignment. The red box indicates the original position of the image before affine transformation. **d**, Stacking of the unaligned coordinates of all slices on the same coordinate system, as well as the aligned coordinates using MOSAICField physical alignment, STalign, PASTE, SANTO, and CAST. **e**, Jaccard Similarity between each pair of adjacent aligned slices from each method. **f**, Unimodal and multimodal feature correlations between adjacent slices computed under MOSAICField physical alignment, MOSAICField affine alignment, STalign, and unaligned coordinates.

For physical alignment, we apply MOSAICField on a global scale. For each pair of adjacent slices, MOSAICField first computes an affine transformation **T** to capture global translation, rotation, and scaling, and then refines it into a nonlinear physical alignment *ϕ*_*P*_ (Supplement §S5). All pairwise alignments **T** followed by *ϕ*_*P*_ are composed to register all slices into a common coordinate system. Compared to the raw, unaligned slices (Fig. 3b), the affine transformations estimated by MOSAICField correct for global displacements and scale differences, placing each slice in its correct relative position (Fig. 3c). The subsequent nonlinear transformations further correct residual global distortions and accurately align tissue boundaries across slices, resulting in a coherent, spatially consistent reconstruction of the tissue that is visually superior to the alignments obtained by STalign, PASTE, SANTO, and CAST (Fig. 3d).

We quantify the alignment quality of each method by computing the Jaccard Similarity between each pair of adjacent slices after alignment, with higher Jaccard Similarity indicating a larger overlap area, and thus better quality registration. MOSAICField achieves the highest Jaccard Similarity (*J =* 0.98 to *J =* 1.00 for all pairs) among all the methods (Fig. 3e, Supplement Fig. S4). STalign is the second best performing method with Jaccard Similarity similar to MOSAICField for most of the pairs (*J =* 0.96 to *J =* 0.99) except Z100 - Z135 (*J =* 0.92), which is the farthest pair in the dataset and has substantial differences in the tissue geometry (Supplement Fig. S3). The other methods output alignments with substantially lower overlap between aligned pairs (PASTE *J* = 0.85 to *J =* 0.96, SANTO *J* = 0.73 to *J* = 0.92, CAST *J* = 0.16 to *J* = 0.95).

While Jaccard similarity measures rough overlap between shapes, feature correspondence provides more fine-grained analysis for alignment quality. Thus, we further assess alignment quality by examining the correlations between gene and protein expression at aligned locations across adjacent slices. Specifically, we compute the average Pearson correlation between protein expression in Z195 and Z210 for all 25 proteins in the CODEX panel, as well as the average Pearson correlation between protein expression in Z210 and gene expression in Z215 for 19 gene–protein pairs whose corresponding coding genes are present and have sufficient signal in the Xenium panel (the remaining six proteins do not have an encoding gene in the Xenium panel; Supplement §S5). More accurate alignments yield higher correlations between protein–protein and gene–protein pairs at aligned locations. As baselines, we calculate correlations using the original unaligned slice coordinates and using only affine transformation without MOSAICField’s nonlinear physical alignment. MOSAICField’s physical alignment substantially increases both unimodal protein–protein and multimodal gene–protein correlations, achieving on average a tenfold improvement over unaligned baselines and a 1.8-fold improvement over affine alignment (Fig. 3f). While STalign also improves correlations relative to unaligned data, its gains are smaller than those from affine alignment, suggesting that MOSAICField’s affine component already captures global structure effectively, and its nonlinear refinement further corrects residual global distortions. A closer examination of every pair of protein-protein and protein-gene correlations further confirms that MOSAICField infers accurate global physical alignment with high unimodal and cross-modal feature correlations (Supplement Fig. S5, S6). We report MOSAICField physical alignment runtime in § S5.3.

### 3.3 MOSAICField morphological alignment tracks the prostate ductal system in reconstructed 3D multimodal tissue

The intricate arrangement of ducts inside the prostate facilitates the transport of prostatic fluid, and understanding the architecture and molecular characteristics of the ductal system is crucial for studying prostate function and pathology [19, 41]. After physical alignment reconstructs a common coordinate system for the HTAN HT891Z1 sample, we use MOSAICField to perform morphological alignment on a local scale to track ducts across the prostate in 3D. We focus on a tumor region of size 1*mm* × 1*mm* (Fig. 4a) and use MOSAICField to compute a nonlinear morphological alignment *ϕ*_*M*_ on this local region between each adjacent physically aligned pair. We manually annotate four ductal regions of interests (ROIs) on the top slice Z215 (Fig. 4a,d) and evaluate MOSAICField’s ability to track the ductal morphology in 3D. We report MOSAICField morphological alignment runtime in § S5.3.

**Fig. 4:**
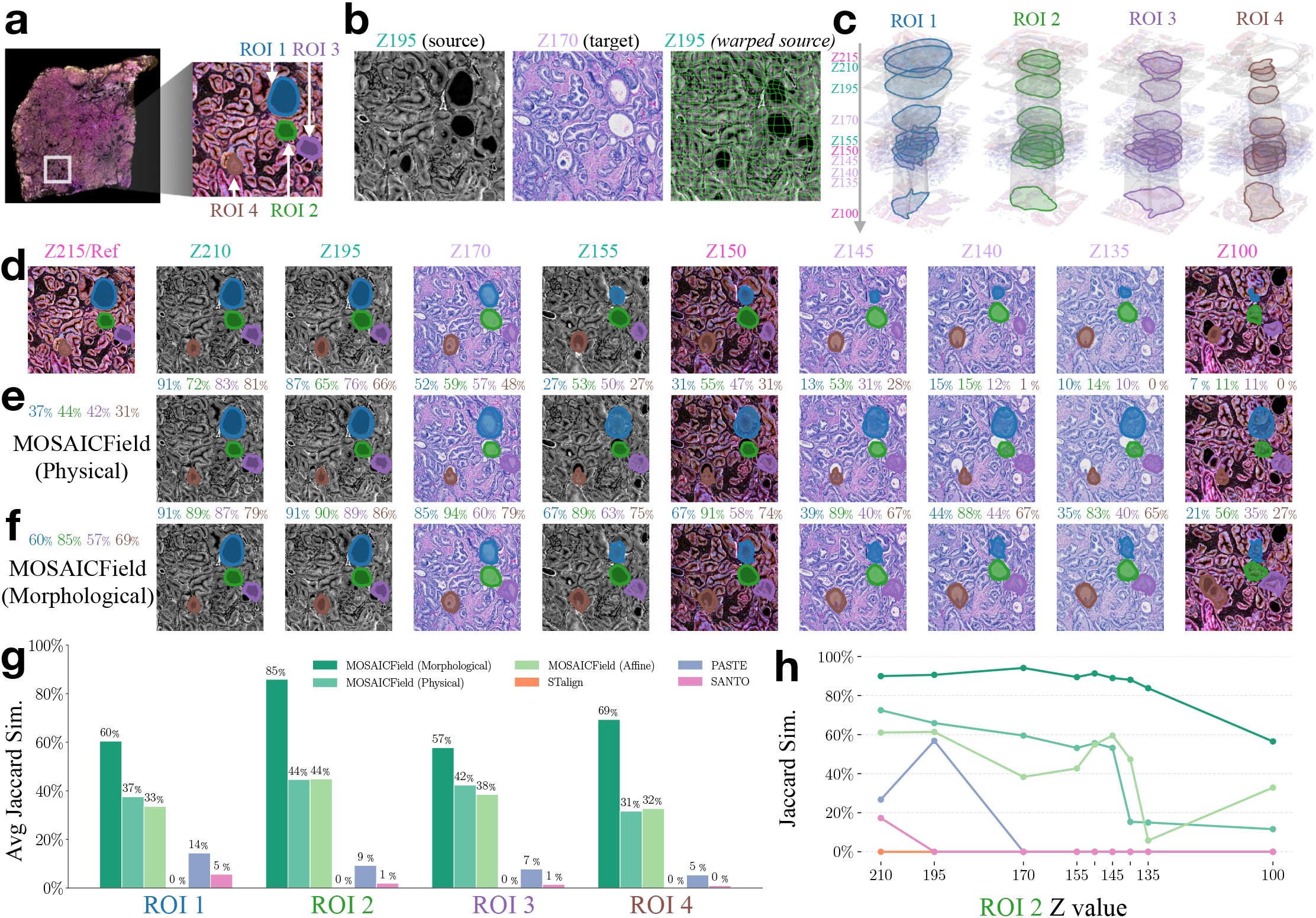
MOSAICField morphological alignment of ductal systems agrees with manual annotation. **a**, A 1mm*1mm local region and four ROIs on Z215. **b**, MOSAICField alignments between Z195 and Z170. (Left, middle) Z195 and Z170 following physical alignment *ϕ*_*P*_. (Right) Z195 after morphological alignment *ϕ*_*M*_ with green curves showing deformation of square grid. **c**, Morphology changes of each ROI tracked by MOSAICField across the physically aligned 3D model. **d**, Manual reference annotations of ROIs across all physically aligned slices. **e**, Tracking of ROIs based on physical alignment only, with Jaccard similarity (reported as percentage) shown above each plot, color coded for each ROI. **f**, Tracking the same set of ROIs using MOSAICField morphological alignment. **g**, Average Jaccard similarity for each ROI with reference annotations on all slices for each benchmarked methods. **h**, For ROI 2 (dark green in panels a,d,e,f), Jaccard similarity with the reference annotation on each slice ordered by decreasing Z.

While physical alignment provides correction against large-scale global deformations between slices, morphological alignment performed locally at full resolution reveals more intricate spatial patterns. For example, ROI1 and ROI2 on the top right part of Z195 (CODEX) have both changed shape between Z195 and adjacent Z170 (HE) (Fig. 4b). The physical alignment (as seen in the “source” and “target” column) roughly aligns the centroid of these ducts, but does not attempt to match their shape. In contrast, MOSAICField morphological alignment (“warped source” column) matches the ducts in the target image both in location and in shape, enabling fine-grained tracking of these morphological features with complex shape changes by learning nonlinear shrinking and expansions (as shown by the green deformation field). Similar observations can be made for other adjacent pairs of slices (Supplement Fig. S7). For each ROI on Z215, we use morphological alignments to track the shape and position down to Z100, where the continuity of ductal structures can be visually validated, and we create a 3D visualization of each duct’s morphology across the stack. The morphological alignment recovers vastly different local deformations for different morphologies (Fig. 4c). ROI1 gradually shrinks and closes, ROI4 expands down the stack, while ROI2 and ROI3 exhibit predominantly cylindrical architectures but with significant shape distortions.

To validate the accuracy of MOSAICField’s morphology mapping, we manually annotate the corresponding ROIs on the entire stack and compare the morphology tracked by MOSAICField with the manual annotation. We visualize the annotated ducts (Fig. 4d), shape and position of ducts tracked by MOSAICField’s physical alignment (Fig. 4e), and by MOSAICField’s morphological alignment (Fig. 4f). We calculate the Jaccard similarity between MOSAICField-tracked duct and manual annotation on each slice, and find that MOSAICField morphological alignment further improves over physical alignment in terms of morphology tracking (see percentage score above the images in Fig. 4ef).

MOSAICField morphological alignment also substantially outperforms STalign, PASTE, and SANTO (Fig. 4g-h, Supplement Fig. S8-S10; We exclude CAST from this comparison given its poor performance on earlier benchmarks). Using the average Jaccard similarity across all slices for each ROI as a measure, MOSAICField has a relative improvement of 35% to 100% compared to the second best alignment method (Fig. 4g). Further examination of the Jaccard similarity score on ROI2 across all Z values shows that MOSAICField hold a consistent advantage over competing methods across all ranges of Z (Fig. 4h; Supplement Fig. S10 for other ROIs). PASTE and SANTO fail to track local morphologies across multimodal slices because they are both designed for unimodal scenarios and they both compute rigid transformations. STalign also fails to compute reasonable morphological alignments possibly due to its reliance on global tissue geometry and variation in cell densities which are not strong signals in local regions.

### 3.4 MOSAICField morphological alignment infers multimodal functional relationships

In addition to comparing with manual annotations, we also evaluate the pixel-level feature correspondence between unimodal and multimodal slices to further validate the morphological alignment computed by MOSAICField. As discussed above, we extract features from Xenium (spatial transcriptomics) by individual gene expression, and extract features from CODEX (spatial proteomics) by individual antibody expression. A well calibrated morphological alignment should see strong correlation between the same features from unimodal slices, and between functionally linked features from cross-modal slices at aligned locations.

As a visual example, we examine the spatial expression of protein CD44 on CODEX slice Z210, which should be strongly correlated with the encoding gene *CD44* on Xenium slice Z150, especially within the spatial region of annotated ductal ROIs (Fig. 5a). We find that with Z210 and Z150 physically aligned, while the ductal ROIs are roughly matched, the *CD44* gene expression on Z150 shows only a week visual agreement with the CD44 protein expression on Z210 in ROIs. In contrast, MOSAICField’s morphological alignment accurately tracks Z210’s ductal ROIs to their corresponding locations on Z150, where the *CD44* gene expression in the tracked ROIs exhibits a strong correlation with the CD44 protein expression on Z210.

**Fig. 5:**
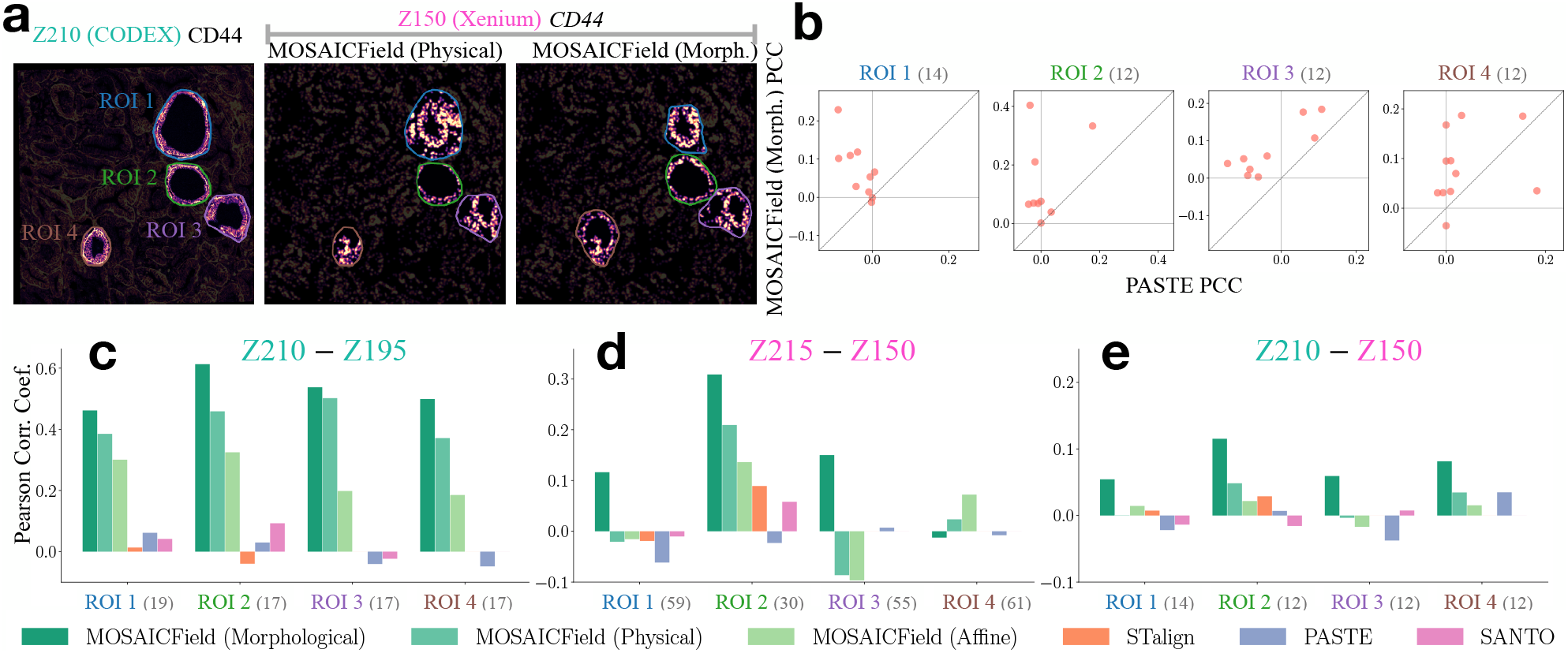
MOSAICField morphological alignment achieves high pixel-level correlations across modalities. **a**, Morphological alignment on a pair of cross-modality slices. (Left) Spatial pattern of protein CD44 within each ROI on Z210. (Middle) Spatial pattern of gene *CD44* on Z150 within each ROI without morphological alignment. (Right) Same as middle but morphologically aligned by MOSAICField. **b**, For this pair and for each ROI, comparing the pixel-level Pearson Correlation Coefficient (PCC) of MOSAICField (Y-axis) to PASTE (X-axis). Each dot represents a gene-protein pair with sufficient expression. **c**, For the CODEX unimodal pair (Z210 and Z195) and for each ROI, average pixel-level PCC for all sufficiently expressed proteins for each benchmarked methods. Number in parentheses following each ROI name is number of features with sufficient expression and counted towards the average calculation. Out-of-boundary mappings are given a correlation score of zero. **d**, Similar to **(c)**, but for the Xenium unimodal pair (Z215 and Z150). **e**, Similar to **(c)**, but for the cross-modal pair (Z210 and Z150).

On three representative pair of slices with paired features, MOSAICField morphological alignment consistently achieves the highest pixel-level average Pearson correlation between paired features among all methods in every ROI (Two CODEX slices in Fig. 5c, two Xenium slices in Fig. 5d, and a CODEX-Xenium pair in Fig. 5e; also see Supplement §S5.2 for feature filtering based on coverage), except ROI4 between Z215 and Z210 (Fig. 5c) because the ductal structure of ROI4 does not exist in Z215 (Supplement Fig. S7). For a closer inspection, we plot the per-feature correlation of MOSAICField morphological alignment against PASTE on the cross-modal pair (Fig. 5b; Supplement Fig. S11 for other pairs and methods), and observe that the morphological alignment step improves the feature correlation across the genes and proteins.

Additionally, we study the generalizability of MOSAICField by holding out features in cross-modal and unimodal alignments. Specifically, we run MOSAICField morphological alignment on the stack while holding out gene *CD44* in Xenium slices and protein CD44 in CODEX slices, and visualize the spatial pattern of the holdout gene *CD44* on Z150 within each ROI morphologically aligned by MOSAICField. Even with CD44 held out, MOSAICField still tracks its correct spatial distribution (Fig. S9a). We then run MOSAICField while holding out a random subset of 90% of the genes, and compute average pixel-level PCC for the holdout genes at aligned pixels. We find MOSAICField achieves superior performance on the holdout genes compared to competing methods even when thoss genes are not used in computing the alignment (Fig. S9b).

## 4 Discussion

We present MOSAICField, a method that aligns spatial slices from arbitrary combinations of experimental modalities using a novel formulation of global invariant optimal transport [1, 16] and neural deformation fields [6, 55] with multi-channel, multi-modality data. We also propose the locally consistent transport problem (Supplement §3) as a unifying theoretical framework for the multimodal alignment problem. In contrast to existing spatial alignment methods [11, 25, 29, 32, 34, 56, 65, 69], MOSAICField distinguishes two distinct alignment formulations: physical alignment and morphological alignment. The physical alignment registers all slices onto a common coordinate system to reconstruct the 3D tissue from 2D measurements, aiming to recover the shape and organization of the original tissue *in silico*. The morphological alignment aims to track morphological structures such as ducts, blood veins, or tumors within the 3D tissue.

MOSAICField offers a unified framework for both alignment problems on diverse experimental modalities, opening up exciting possibilities for downstream spatial research that were previously challenging or unattainable. We previewed these possibilities by using MOSAICField to build a multimodal 3D model of a prostate tumor by aligning spatial slices from transcriptomics, multiplex imaging, and histological staining. The construction of accurate multimodal 3D tissue models is a key goal of collaborative efforts to build comprehensive 3D tissue and tumor atlases [22, 39, 42, 50] which will enhance our understanding of tissue heterogeneity, disease progression, and cellular microenvironments in 3D [42]. The ability to track morphological features such as ductal systems or tumor boundaries will facilitate the study of dynamic biological processes, such as tumor invasion or angiogenesis [64].

There are multiple opportunities to extend MOSAICField. First, we distinguish physical versus morphological alignment by applying MOSAICField on two different scales. There are other options for making this distinction, such as to define the two problems through different rigorous mathematical formulations. Another promising avenue is to extend MOSAICField to support time-series spatiotemporal data, enabling the study of dynamic biological processes [17, 29], such as wound healing, immune response, or tissue regeneration. Third, our algorithm shares fundamental mathematical limitations with a diffeomorphism: it does not model the creation of new morphologies that lack corresponding support in the source slice. To our knowledge, existing registration approaches face the same challenge, and addressing this limitation is a promising future direction. Finally, integrating MOSAICField with downstream computational models for tasks like cell-cell communications [24] or multimodal feature generation [31] could open up new possibilities for hypothesis generation and validation in spatial biology.

## 5 Code Availability

Source code of MOSAICField is available at https://github.com/raphael-group/MOSAICField.

## 6 Acknowledgments

We thank the patients, staff and scientists who contributed to this study as well as the NCI, Human Tumor Atlas Network (HTAN) consortium and Genomic Data Analysis Network (GDAN). We thank Washington University Center for Cellular Imaging. We thank Hirak Sarkar, Wen-hung Chou, Feng Chen, Clara Liu, Chia-Kuei Mo, André Targino for their feedback throughout the project.

This research was supported by NIH/NCI grant U24CA248453 to B.J.R. and U2CCA233303 to L.D. J.G. gratefully acknowledges support from the Schmidt DataX Fund at Princeton University made possible through a major gift from the Schmidt Futures Foundation.

## Supplementary Material

### S1 Solving the affine alignment problem

We set *c*_1_ and *c*_2_ to be Euclidean distance in a lower-dimensional feature space. Here we use Principal Component Analysis (PCA) to reduce the dimension of the feature space, but nonlinear dimensionality reduction techniques such as *k*–Nearest Neighbor approximation [7, 14] or autoencoders [36] can also be used. Since costs *c*_1_ and *c*_2_ might be on a different scale for different modalities, we scale the costs such that 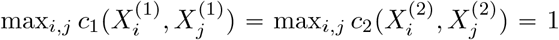. **T** : ℝ^2^ → ℝ^2^ is represented by a transformation matrix **T** ∈ ℝ^3×3^ for 2-dimensional points in homogeneous coordinates.

#### Optimization

We solve for both **T** and **Π** simultaneously. We employ an alternating optimization between **T** and **Π**, in the flavor of block coordinate descent, on the “parameter” **T** and “posterior” **Π**. When **Π** is fixed, the optimization problem for **T** becomes the following:

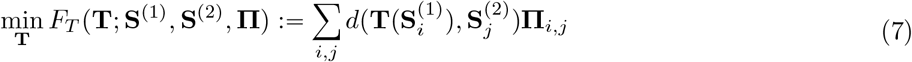

This becomes a least squares regression when we require **T** to be an affine transformation (or a Procrustes problem [59] when we require **T** to be a rigid transformation). On the other hand, when **T** is fixed, the optimization problem for **Π** can be reorganized as follows:

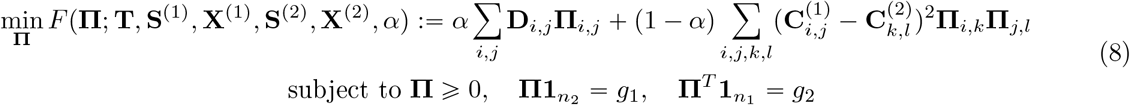

Here, we define the (now constant) matrices 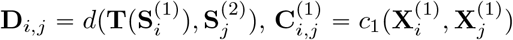, and similarly 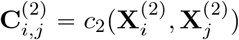. This is an instance of Fused Gromov-Wasserstein Optimal Transport, which can be solved by an iterative conditional gradient approach proposed in [57].

The training procedure is in Algorithm 1, where *K* is the number of alternating steps. We let function FGW-OT(**D, C**^(1)^, **C**^(2)^, *α*) be the call to the Fused Gromov-Wasserstein Optimal Transport solver, and Projection(**S**^(1)^, **S**^(2)^, **Π**) be the linear regression solver for affine transformations.

##### Algorithm 1

Affine alignment step of MOSAICField

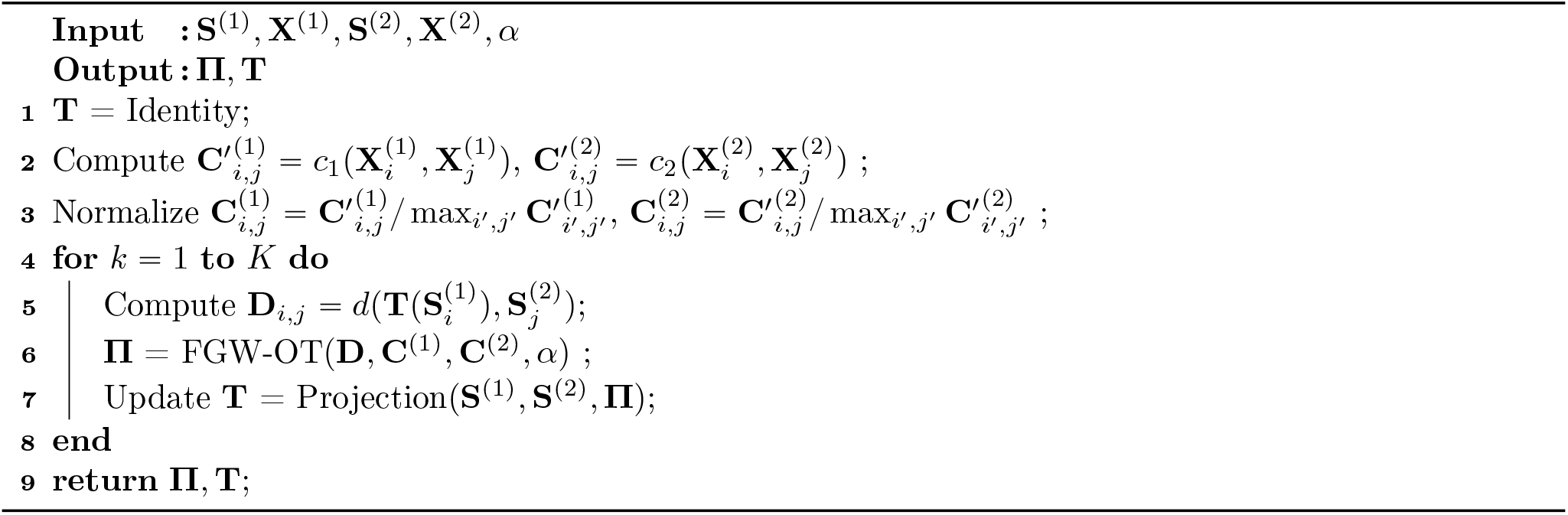

### S2 Optimization for Nonlinear Transformation

*Regularization on ϕ* The regularization consists of two terms

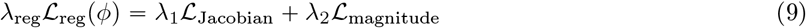

ℒ_Jacobian_ penalizes negative Jacobian determinants of *ϕ* and is given by

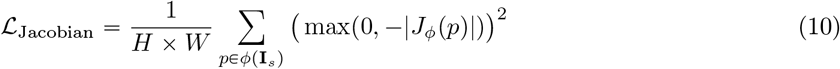

where *J*_*ϕ*_(*p*) is the Jacobian matrix of *ϕ* at position *p*. This term encourages the deformation field *ϕ* to preserve local orientations. Following [63], we implement *J*_*ϕ*_(*p*) using local finite difference approximation. The second term, ℒ_magnitude_, regularizes the magnitude of the displacement **v**_*θ*_ and is defined as:

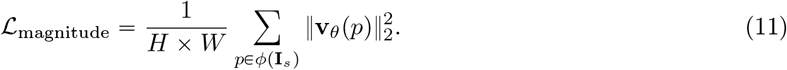

*Optimizing for ϕ* Given the images **I**_*s*_ and **I**_*t*_, we optimize *ϕ* using gradient descent on the combined loss function (Equation 2). We use spatial transformer networks [23] to compute gradient flow through spatial transformations.

### S3 Locally Consistent Transport Problem

In this section, we propose the concept of Locally Consistent Transport Problem, which is an extension of Gromov-Wasserstein optimal transport, and is tightly connected to how our neighborhood similarity loss (referred to as MIND loss in this section) is implemented. By the end of Section §S3.4, we will propose a new formulation of multimodal loss function that is flexible with strong connections to both MIND-like losses and Gromov-Wasserstein optimal transport.

#### S3.1 Definition

Let (𝒳, *k*_𝒳_, *µ*_𝒳_), (𝒴, *k*_𝒴_, *µ*_𝒴_) be two metric-measure spaces, where (*k*_𝒳_, *k*_𝒴_) are distances while (*µ*_𝒳_ ∈ 𝒫 (𝒳), *µ*_𝒴_ ∈ 𝒫 (𝒳)) are measures on their respective spaces. Let *T* : 𝒳 → 𝒴 be a Monge transport map transporting *µ*_𝒳_ to *µ*_𝒴_, i.e.

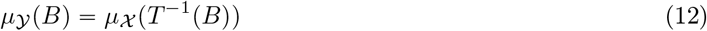

for all *µ*_𝒴_-measurable sets *B*. We also assume an additional metric *k*_*S*_ over 𝒳. Given *ϵ >* 0, we define the *locally consistent transport problem* as minimizing

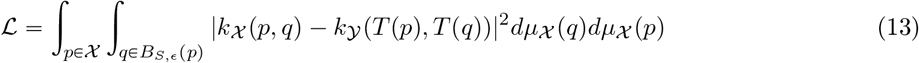

over *T*. *B*_*S,ϵ*_(·) is the *ϵ*-ball with respect to metric *k*_*S*_ (meaning that *k*_*S*_(*p, q*) ≤ *ϵ*).

#### S3.2 Connection to Gromov-Wasserstein

We prove that Gromov-Wasserstein OT (in absence of the third metric *k*_*S*_) is a specific instance of our proposed problem.

##### Proposition 1.

*At ϵ* → ∞, *the locally consistent transport problem becomes the Gromov-Wasserstein optimal transport over Monge maps*.

*Proof*. Recall the Gromov-Wasserstein optimal transport problem with coupling *π* (remember *T* pushes *µ*_𝒳_ to *µ*_𝒴_):

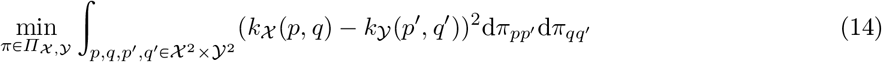

Now we define the natural coupling from the transport map *T* by a push-forward operator:

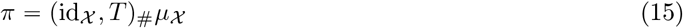

In the discrete case, this means *π*_*i,T* (*i*)_ *µ*_𝒳_ (*i*) and *π =* 0 elsewhere. Substituting this back to the GW formulation, we have:

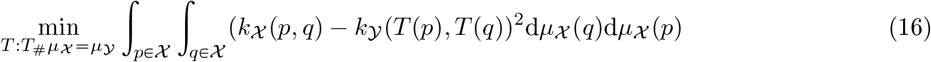

This gets us the formal equivalence with equation 13, with *B*_*S,ϵ*_(*p*) = 𝒳 or equivalently *ϵ* → ∞.

#### S3.3 Connection to Current Implementation: No Transformation Scenario

In this section and the following section, we establish the connection of our proposed locally consistent transport problem to the MIND formula. For this section, we start by specifying the loss function in a more general manner than the main text. This is followed by two ways to establish equivalence to locally consistent transport problem, although both with some caveats.

Let *I* be an image with discrete pixels: *I* ∈ ℝ^*H*×*W* ×*C*^ where *C* denote the number of channels in *I*. Given hyperparameter *ρ*, we define:

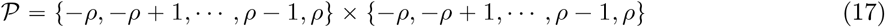

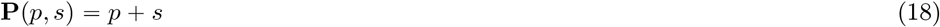

as the patch offsets and patch function. Given two pixel locations *p* and *q*, we define a patch descriptor 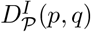 as:

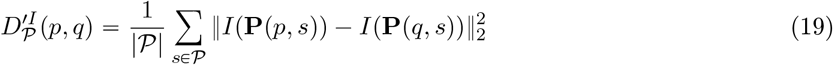

Next, we are given a second hyperparameter *r*, and from *r* we define

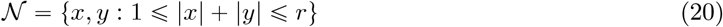

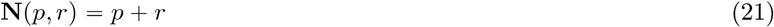

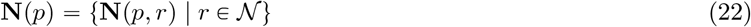

as the neighborhood offsets, the neighborhood function and neighborhood set. Given image *I*, we first define the average patch over neighborhood:

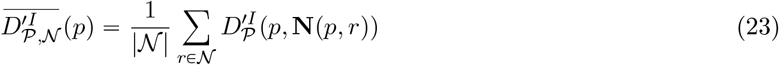

From this we define the neighborhood descriptor, which is a transformed and normalized version of the average patch for *q* ∈**N;**(*p*):

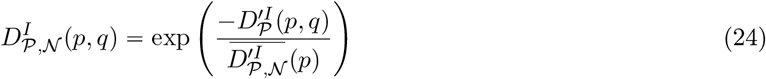

And finally given two images *I* and *J* of exact same size, the MIND loss between them are defined as:

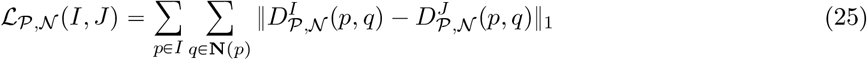

Now assume we train an transformation *ϕ* from image *I* to *J*, and use *ϕ*(*I*) to denote the post-transformation image. Currently, we uses ℒ _𝒫, 𝒩_ (*ϕ* (*I*), *J*) as the loss function. We show two ways of fitting this into our current framework. The first way is by not using *T* in the optimal transport formulation, and directly setting 𝒴 = *ϕ*(*I*):

##### Proposition 2.

*The MIND loss is a specific instantiation of locally consistent transport problem between images 𝒳* = *J and 𝒴* = *ϕ*(*I*), *with trivial transformation T* = *id*_𝒳_, *µ*_𝒳_ *being a uniform distribution over pixels of J (similarly for µ*_𝒴_ *and* 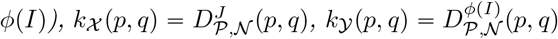 *and k*_*S*_(*p, q*) = ‖*p* − *q*‖_1_ *leading to B*_*S,ϵ*_(*p*) = 𝒩 (*p*) *with hyperparameter r* = *ϵ*

*Proof*. Plugging everything into Equation 13, recalling that in this setup we have *T p* (*p*), and replacing integrations with summations now we are operating over discrete space we have:

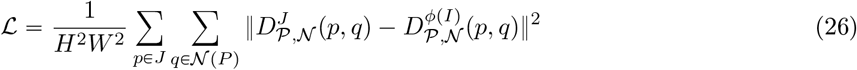

which is the main loss function but with *ℓ*_2_ norm instead.

Alternatively, we can try to connect *T* and *ϕ*. The side effect of doing this is to introduce an extra dependency of *T* over *k*_𝒴_ :

##### Proposition 3.

*Given I, J, ϕ, there exists a formulation of locally consistent optimal transport with 𝒳 = J, 𝒴* = *I, T* = *ϕ*^−1^, *and a definition of k*_𝒴_ *which depends on T, such that it also instantiates the MIND loss*.

*Proof*. We can simply define

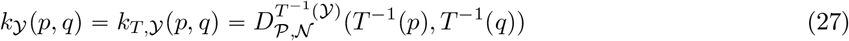

and *k*_𝒳_ (*p, q*) and *k*_*S*_ (*p, q*) following the previous proof. Plugging these values into Equation 13 again and using the same procedures as last proposition, we have:

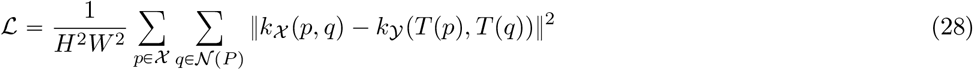

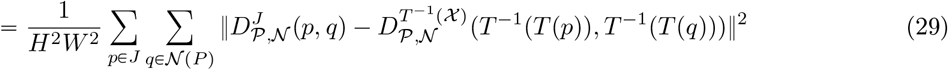

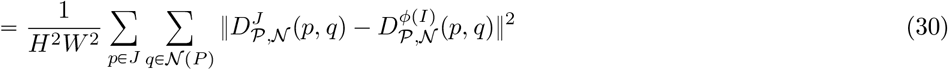

At which point we come back to Equation 26. Alternatively, we observe that the only reason this dependency is necessary is because *ϕ* distorts the definition of neighborhoods and patches. For this reason, we can write down an alternative formulation where we redefine the neighborhood and patch functions **P, N** (recall that the original definitions are **P**(*p, s*) = *p* + *s*, **N**(*p, r*) = *p* + *r* and *T* = *ϕ*^−1^):

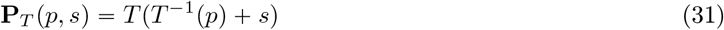

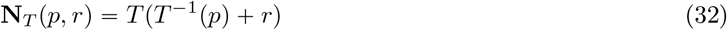

From here we can redefine every other term up to *k*_*T*,𝒴_ and arrive at the same conclusion. We derive the exact constructions and correspondences in Supplemental Section §S3.5.

#### S3.4 Context-free MIND

We now propose a modified problem that strictly fits within our proposed framework but is no longer strictly equivalent as follows.

- *𝒳* and *𝒴* are the discrete set of pixels in image *J* and *I*, and *T* = *ϕ*^−1^.
- 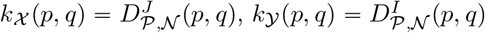

Now *k*_𝒴_ is no longer dependent on *T*. We can plug in Equation 13, using the similar procedures as last proof we have:

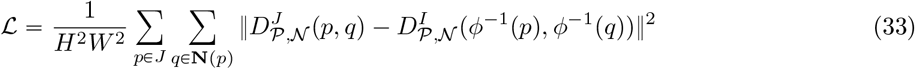

The terms involving *J* are already matched to Equation 26. However the following equation does not hold in general:

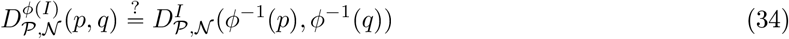

unless **N**_*ϕ*_ (*p, r*) **= N** (*p, r*) (and similarly for **P**), meaning *ϕ* is a simple translation. In other words as long as *ϕ* changes geometry we lose the strict equivalence. Fig. S1 give a more visual example of *ϕ* changing geometry leading to different **N**.

**Fig. S1:**
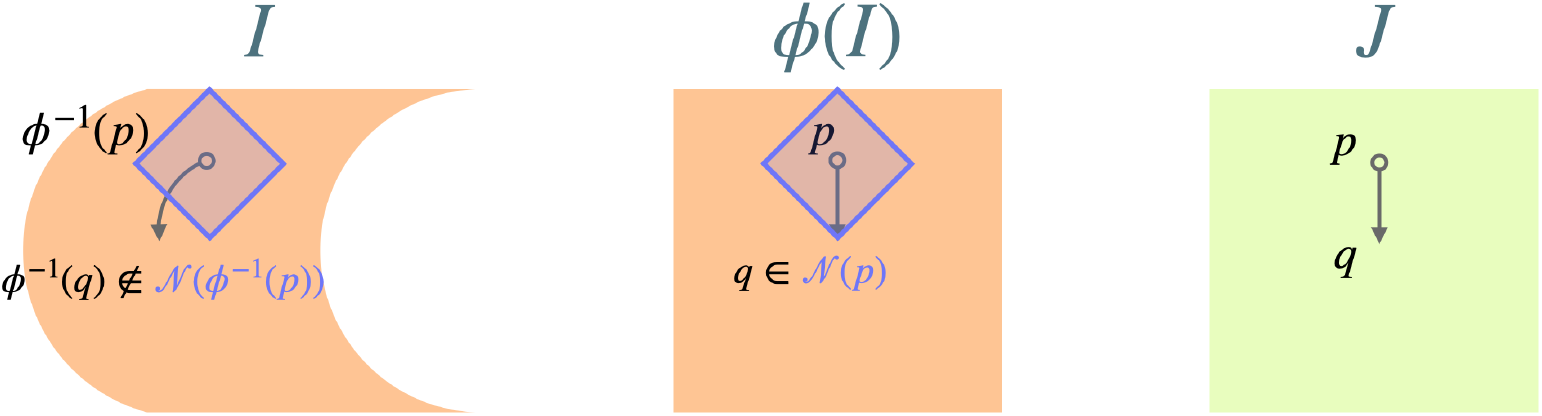
A hypothetical scenario showing why the context-free MIND is not strictly equivalent to our current loss function. In this version, *I* is a squeezed image and *ϕ* stretches it back to square like *J*. Given a pixel *p* and *q* ∈ **N**(*p*) on *ϕ*(*I*), we can see the relation no longer hold on *I* after we apply *ϕ*^−1^ to pixel coordinates.

For this manuscript, we opt with implementing the generalized MIND as described in the main text. We plan to implement context-free MIND (as a specific instantiation of locally consistent transport problem) in the future, and explore more variants based on this formulation as well.

#### S3.5 Addendum: Reconstructing MIND loss with uneven geometry

Here we fill in the missing constructions in the second proof to Proposition 3. For the sake of convenience we switch back to using *ϕ* in these constructions. We start with the following

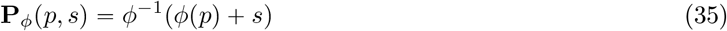

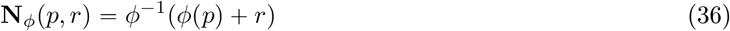

and from here:

With these changes we can further define 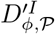 and 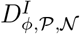:

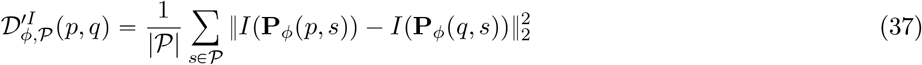

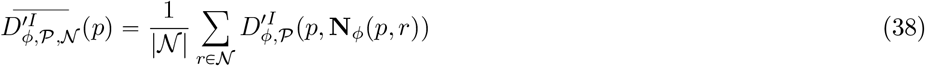

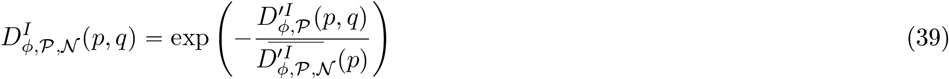

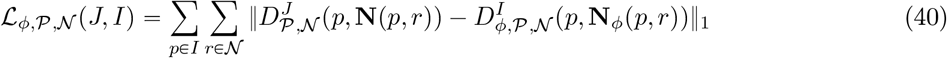

Equations 37,38,39,40 ensures the following equality always hold with respect to Equations 19,23,24,25 in the following way:

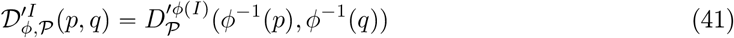

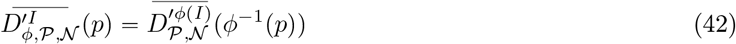

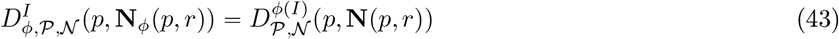

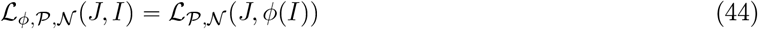

And at this stage we can formally set

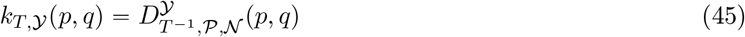

(parameter change from Equation 39), and the loss function would be

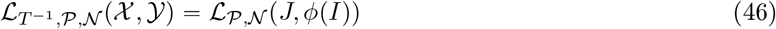

at this point we come back to the original MIND loss formulation and finish the proposition proof. In other words, by only changing the definition of **N** and **P**, we can ensure all computations are performed on the transformed image *ϕ*(*I*).

### S4 Generation and evaluation of simulated data

#### S4.1 Physical alignment simulation

We use the Stereo-seq E10.5 mouse embryo slice from [9] as the reference target slice. This slice contains 18408 cells with 25201 genes measured. We use the original spatial coordinates of the slice as the target slice, and generate a globally distorted source slice in the following way. Let 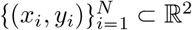 be the original point cloud. Without loss of generality assume they are centered with centroid (0, 0). Let *L*_*x*_ max (*x*) min (*x)* be the horizontal span (range in *x*). Let *L*_*y*_ = max(*y*) − min(*y*) be the vertical span (range in *y*). For each point (*x, y*) on the original slice, we map it to

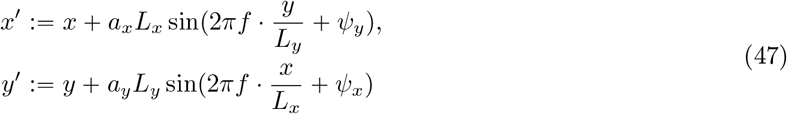

where *a*_*x*_, *a*_*y*_ are amplitude parameters and *f* is a frequency parameter denoting the number of oscillation cycles across the full span. We set *a*_*x*_ *a*_*y*_ 0.03 and *f* 3 for our simulation. *ψ*_*x*_, *ψ*_*y*_ are random phase offsets in the sine waves so the ripple does not align to axes too neatly

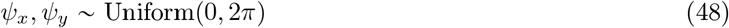

This pattern adds a smooth, nonlinear distortion to the original shape but without tearing or folding. After the sine warp, the point cloud coordinates may become a bit larger than the original shape, so we scale the *x* and *y* span of the warped point cloud to the original spans to keep the overall size of the shape. We then scale the entire slice by 1.1 × and rotate it clockwise by 25 degrees to simulate tissue scaling and translation that commonly happens during sample preparation. The spatial coordinates of cells after this nonlinear warping become the source slice, where our goal is to use spatial alignment methods to align the source slice onto the target slice to correct for the global distortion.

Since our method aligns slices with distinct modalities, we need to simulate features such that the source and the target slices come from different modalities. To simulate multimodal features, we divide the gene set of 25201 genes into two halves, each with 12600 genes. We assign one gene set to the source slice and the other to the target (that is, the gene expressions of cells in each slice only contain half of the total genes measured), so that the two slices do not share any common gene. We then use Principle Component Analysis (PCA) to reduce the feature dimension of each slice from 12600 to 50, and multiply the feature matrix **N** of the source slice by a random matrix **X**_*new*_ **X**_*old*_**W**, where **W** ∈ ℝ^50×50^ is a random matrix with entries drawn from a standard normal distribution. This operation applies a random change of basis to the source slice’s feature space, still preserving linear relationships between features but could cause trouble for alignment methods that rely on strict feature matching.

#### S4.2 Morphological alignment simulation

To simulate a morphological alignment problem, we begin with the same E10.5 embryo slice. For the target slice, we embed a hollow circular region within the tissue to represent a luminal structure. To create the source slice, we embed the lumen in a slightly shifted position such that the two lumens partially overlap, simulating a scenario where a morphological structure traverses the tissue at an oblique angle relative to the slicing direction. We generate multimodal features for both slices following the same procedure described above. Additionally, because cells surrounding a morphological structure in real data often share similar molecular characteristics, we assign zero-valued 50-dimensional features to a ring of cells surrounding the lumen in both slices.

We rasterize the source and target slice point clouds into multi-channel images to run MOSAICField on both simulations, where the number of channels is the number of features.

#### S4.3 Evaluation metrics

##### Registration Error

We use Registration Error to evaluate the accuracy of alignments for the physical alignment simulation. In the physical alignment simulation, since the source slice coordinates are a nonlinear transformation of the target slice coordinates, we know the ground truth correspondence between cells in the source and target slices. Then, the registration error is defined as the average distance between each cell on the source slice after alignment and its corresponding cell on the target slice

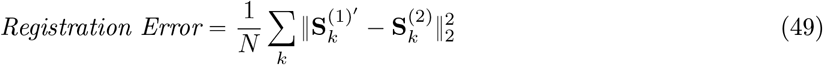

where 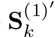 and 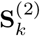 denote the coordinate of the *k*-th cell on the aligned source and target slice, respectively. A smaller registration error indicates that each cell on the source slice is close to its ground truth corresponding cell on the target slice, hence a more accurate alignment.

##### Label Transfer Adjusted Rand Index (LTARI)

LTARI, defined in [34], measures how the alignment matches cells of the same type across slices. Specifically, for each cell on the target slice, we assign it a new cell type label by assigning the cell type label of the cell closest to it on the aligned source slice. That is, we define the new cell type label of each cell on the target slice to be the cell type label of the cell closest to it on the source slice after alignment. This assignment transfers the cell type annotation from the source slice to the target slice. Then, we compare this transferred labeling with the ground truth labeling of the target slice, and compute the ARI of the two clusterings. A high LTARI indicates that the alignment maps each cell to some cell on the other slice with the same cell type label, hence a better alignment.

##### Jaccard Similarity

To evaluate the alignments in morphological alignment simulation, and later in real data, one of the metrics we use is the Jaccard Similarity between two point clouds. We define the Jaccard Similarity between two point clouds *P, Q* in the following way. For each *p* ∈ *P* and *q* ∈ *Q*, define *ϵ*-neighborhoods

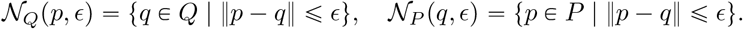

The intersection set is the collection of all unique points from *P* and *Q* that are within *ϵ*-distance of at least one point in the opposite point cloud:

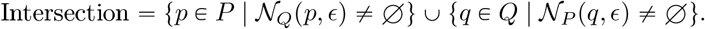

The union set consists of all unique points in *P* and *Q*:

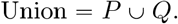

The Jaccard Similarity is given by the classical intersection over union:

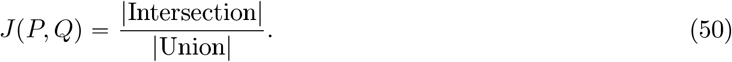

A high Jaccard Similarity indicates a better alignment because of a larger overlap in point cloud coordinates. We use *ϵ =* 1 for the morphological alignment simulation dataset as cells in Stereo-seq are on a pixel coordinate space.

#### S4.4 Ablation studies

We conduct ablation studies to establish the contributions of various components in the multimodal objective function to the performance of MOSAICField. Specifically, we apply MOSAICField to the physical alignment simulation above with varying *β* (Eq. 6), *λ*_1_, *λ*_2_ (Supplement Eq. 9) to establish the contributions of the MIND loss, the MSE loss, and the regularizations of the MOSAICField objective function. *β* ≈ 0.5 achieves the best performance in terms of both Registration Error and LTARI for the simulated multimodality pair, indicating that both the multimodal MIND loss and the single-channel MSE loss are essential for the performance of MOSAICField (Fig. S2a). To test whether the single-channel MSE loss would be more beneficial in unimodal alignments than multimodal alignments, we generate another simulated pair of slices according to the procedure described in § S4.1 but without multiplying the feature matrix with a random matrix **W**, so that the features across slices belong to the same modality. Surprisingly, we find that the multimodal MIND loss is still beneficial in the unimodal case (Fig. S2b), achieving better alignments than using MSE loss alone (*β =* 1), probably because the MIND loss optimizes for structural information in addition to feature information. Therefore, we use *β =* 0.5 for all experiments in this work.

We then investigate the effects of *λ*_1_, the strength for the Jacobian regularization, and *λ*_2_, the strength for the magnitude regularization (§ S2). We test *λ*_1_ ∈ {0.0, 2.5, 10.0, 25.0, 50.0} and *λ*_2_ ∈ {0.0, 5 × 10^−6^, 5 × 10^−5^, 0.0005, 0.005} and find that MOSAICField is robust to a wide range of parameter settings (Fig. S2cd). We use *λ*_1_ = 2.5, *λ*_2_ = 5 × 10^−6^ for all experiments in this work.

### S5 Processing and evaluation of HTAN prostate cancer multimodal data

We now detail the steps to find a physical alignment over full slices, and a morphological alignment in a region of interest. We also detail our evaluation procedures.

#### S5.1 Physical alignment

##### MOSAICField setup

MOSAICField affine alignment step takes each spatial slice in the form (**X, S**), where **X** is a *n* by *m* feature matrix and **S** is a *n* by 2 coordinate matrix for *n* locations on a slice. The HTAN prostate cancer data comes in the form of whole-slide images. Whole-slide images are high-resolution, leading to a huge number of pixels such that it is unrealistic to treat each pixel as a location in (**X, S**) individually unless one either scale down the image or uses a low-rank approximation [18]. For example, H&E images can be up to 20,000 by 20,000 pixels in size, consisting of a total of 400 million pixels. Not all pixels are needed for computing the affine transformation in step 1 of MOSAICField. Therefore, a preprocessing step is needed to encode imaging data into a form that can be handled by the MOSAICField affine alignment algorithm, while the resulting transformation can still be applied to all pixels in original resolution.

We assume the full resolution of each image to be one *µ*m per pixel (if not, we first scale the images into this resolution and take this resolution as full-resolution), resulting in an image **I** ∈ ℝ^*C*×*H*×*W*^, where (*H, W*) is the size of the input image, and *C* is the number of channels. We divide **I** into patches of 64 × 64 *µ*m (64 × 64 pixels), and use the Otsu algorithm [46] to select those that contain tissue, excluding the background. We train a convolutional variational autoencoder (VAE) [27] on the set of *C* × 64 × 64 patches and take the latent embedding *z* for each patch as the feature for that patch, and take the pixel location of the center pixel of the patch as the spatial coordinate of the patch. Hence an image **x** is represented as a tuple (**X, S**). **X** ∈ ℝ^*N* ×*k*^ and **S** ∈ ℝ^*N* ×2^, where *N* is the number of on-tissue patches, *k* is the dimension of the VAE embedding. We implement the VAE in PyTorch with five hidden layers for the encoder and five hidden layers for the decoder. The encoder hidden layers dimensions are 32, 64, 128, 256, 512, where each layer is a 2D convolutional layer with kernel size 3, stride 2, and padding 1. The decoder layers have the reverse configuration of the encoder layers. We set *k =* 50, and train the VAE using the Adam optimizer [28] for 10 epochs with a batch size of 64 and learning rate of 0.005.

For Xenium, we also divide the coordinate into 64 × 64 *µ*m patches, and for patches with at least one cell inside, we set the feature of that patch to be the normalized total expression of all cells whose geometric center lies within that patch. We then run PCA dimensionality reduction over the normalized expression matrix. The resulting feature matrix **X** is the PCA components of each patch containing at least a cell, and the resulting coordinate matrix **S** is the center coordinates of these patches.

The (**X, S**) preprocessed from each image can then be registered by MOSAICField with other slices for affine alignment. Note that, since the spatial coordinate of each patch is the location of the center pixel, **S** is in the same scale as the original pixel resolution, hence the affine transformation **T** can be applied to transform all pixels in the original image to obtain a registered image at the original resolution (one *µ*m per pixel).

After obtaining affine aligned images **I**_1_, …, **I**_16_ in the original resolution for all 16 slices, we use the second step of MOSAICField to compute a nonlinear deformation field *ϕ*_*P*_ between each pair of images. Each image is of shape *C* × 7227 × 7614 at the full-resolution of 1 *µ*m per pixel, where *C* is the number of channels for each modality. For H&E, we use *C =* 3 for the RGB channels. For CODEX, similarly we use *C =* 25 for all antibodies. For Xenium we have *C =* 476 for all genes, and we build the image by reading the location of each individual transcript and adding a count of one to the pixel containing the location.

To run MOSAICField on a global scale, we first downscale each image 5 × to *C* × 1445 ×1522, then use MOSAICField to compute a nonlinear deformation field between each pair of down-scaled images. The deformation field is then interpolated (our implementation of the deformation field through the spatial transformer networks [23] scales all coordinates between 0 and 1 so it is easy to interpolate) back to the original resolution as the physical alignment *ϕ*_*P*_ between each pair of images in full-resolution. We set hyperparameters *α* = 0.8, *β* = 0.5, *λ*_1_ = 2.5, *λ*_2_ = 0.000005 for all our experiments.

##### Evaluation of physical alignment

To evaluate the quality of the physical alignment of MOSAICField with that of other methods, we first use the Jaccard Similarity between each pair of registered adjacent slices for each method. To do so, we view the slices as point clouds (**X, S**) as above and then use Equation 50 with *ϵ =* 64.

We then compute the Pearson correlation between aligned locations. This is done in multiple steps: We determine the contour of source slice by rasterizing the grayscale image and fitting an alphashape over the set of foreground patches determined by otsu algorithm, collect the set of interior pixels (on the 5 × downscaled image of pre-transformation data) of the fitted polygon denoted as *P*, then for each tested method, we apply the transformation over *P* and get a new set of coordinates *Q*. We use local interpolation to obtain the readout of relevant features for *P* on the source slice, and for *Q* on the target slice. Denote these readouts *X*_*P*_ and *X*_*Q*_, and the Pearson correlation is calculated as 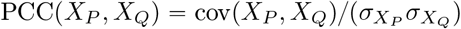.

#### S5.2 Morphological alignment

##### MOSAICField setup

With all slices in the HT891 stack physically aligned to the same coordinate system, we zoom in onto a local tumor region of size 1*mm* × 1*mm* for morphological alignment, corresponding to an image patch of size 1000 × 1000 in full-resolution. As discussed in Method §2.3, morphological alignment assumes the slices are already physical aligned. We use MOSAICField to compute a morphological alignment *ϕ*_*M*_ on this local region between each pair of slices. The *ϕ*_*M*_ between each pair of slices can be composed to track morphologies down the stack. We use the same hyperparameters for MOSAICField as in the physical alignment above.

##### Evaluation of morphological alignment based on ROI annotation

We first generate the data post physical alignment and perform all operation evaluation based on it; Local interpolation is needed to generate the new feature readouts if the readout of a pixel on the post-transformation dataset is not from an integral coordinate in the pre-transformation dataset. We annotate four ductal ROIs on the local region of all relevant slices (Z215 – Z100), manually done with the help of ImageJ as bitmasks. We use the annotation at Z215 as input to all benchmarked methods and hide the rest of annotation for testing.

For our method, we use the Z215 bitmask as input to the displacement field inferred by morphological alignment and gather the pixels with positive intensity at each Z, then fit an alphashape as the inferred polygon. For the affine baseline, we run the bitmask backwards in the global displacement field to obtain its location if we assume it is unchanged under affine coordinate system. For other competing methods, we gather the constituent pixels of the Z215 bitmask, use these methods to compute where these pixels ends up on every other slice, then fit an alphashape as its inferred polygon. We skip the alphashape fitting step if the inferred pixel locations are entirely out of the local region.

To calculate the Jaccard similarity scores as seen in Fig. 4d-f, we calculate the area of union and intersection from reference annotation polygons and inferred polygon by a given method, then set the Jaccard similarity as area of union over area of intersection. This is done for all slices after Z215 with known annotations, and scores over all slices (other than Z215) are averaged per ROI per method to generate Fig. 4e.

##### Evaluation of morphological alignment based on feature mapping

As Xenium is a count-based dataset converted to images, the extreme sparsity of Xenium data becomes an issue. We apply a 2-dimensional Gaussian filter with *σ =* 3 over all features in all Xenium data, ensuring that each transcript have a physical volume, and slight misalignment between transcript readouts still gets credited in the evaluation. CODEX is already microscopy image and thus we opt to not perform such smoothing there.

For the calculation of Pearson correlation, we use a similar approach compared to the last part of physical alignment evaluation with PCC. First, we rasterize the annotation on source slice (post-physical-alignment) into individual pixels (1*µ*m × 1*µ*m sized) as *P*. At this stage, we collect the total count/pixel intensity within the ROI on the source slice. If this number is below 25 (if source slice is Xenium) or 400 (if source slice is CODEX - pixel intensities are normalized to the maximum of 1), we skip this feature pair. The filtering step is necessary because sufficient signal is required for reasonable evaluation, similar to performing quality control in scRNA-seq analyses. For all benchmarked methods, we use them to warp *P* into the coordinate of target slice, denoted as *Q*, and calculate the PCC as done in the previous section. As mentioned in the main text, correlation scores are set at zero outside the ROI boundaries as these indicate alignment failure and should not be simply considered as missing data.

The procedure above involves finding the mapping of a source slice pixel inside our inferred neural displacement field. Since the inferred variable is actually displacement from target slice, we need to numerically invert the displacement field. This is done in two ways.

- First, we run a fixed point iteration: To solve *p* + *ϕ*(*p*) = *q* given *q*, we note *p* is a fixed point of *g* (*p*) *= q* − *ϕ* (*p*). Thus, we start by setting *p*_0_ *q* then iteratively set *p*_*i*_ *= q* − *ϕ* (*p*_*i* −1_) till convergence or after a fixed number of iterations.
- In case the first method does not converge (due to some inferred flows not being locally 1-Lipschitz), a recursive grid search is performed instead. For every point *q* we are trying to solve *p + ϕ* (*p*) *= q* for, we start by dividing the entire image space into 25 equally sized grids. Then, the error is calculated by 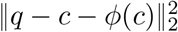 for each grid center *c*, and the 20 grids with highest scoring centroids are retained. Next, we lower grid resolution by 5 ×, split each remaining grid into 25 equally sized subgrids, then repeat the centroid evaluation process and keep the highest 20 scoring grids. This is repeated until the grid size is small enough, or centroid loss is low enough.

#### S5.3 Runtime

For the HTAN HT891Z1 sample, the average runtime for MOSAICField physical alignment for a pair of slices (image size 1445 × 1522) was around 19 minutes on a Nvidia Tesla P100 GPU, while the average runtime for MOSAICField morphological alignment (image size 1000 × 1000) was around 7.5 minutes. We expect the runtime to be much shorter on more modern GPUs.

**Fig. S2:**
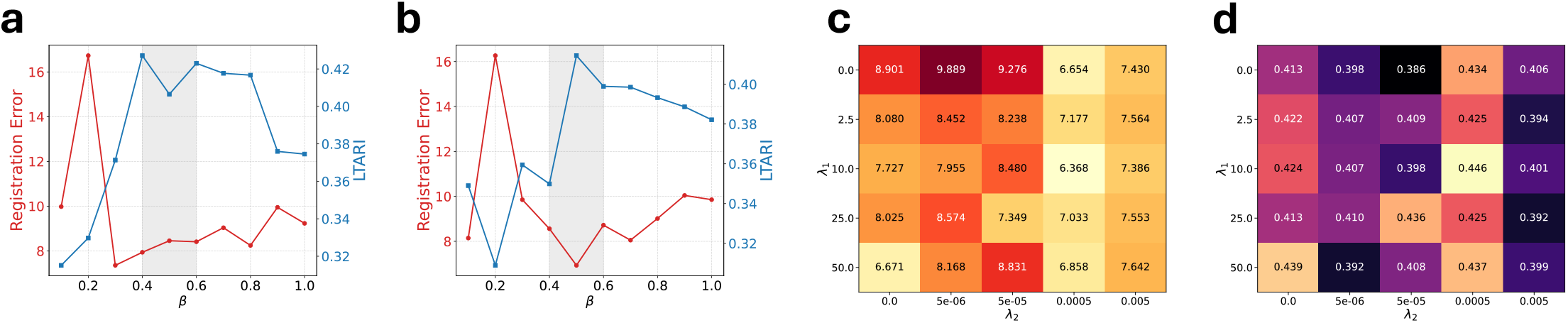
Ablation studies on component attributions. **a**, Ablation on *β* (MIND vs MSE) in Eq. 6 for multimodal alignment measuring Registration Error (lower is better) and LTARI (higher is better). **b**, Ablation on *β* for unimodal alignment measuring Registration Error and LTARI. **c** Registration Error for various parameter combinations of *λ*_1_, *λ*_2_ in Supplement Eq 9. **d**, LTARI for various parameter combinations of *λ*_1_, *λ*_2_.

**Fig. S3:**
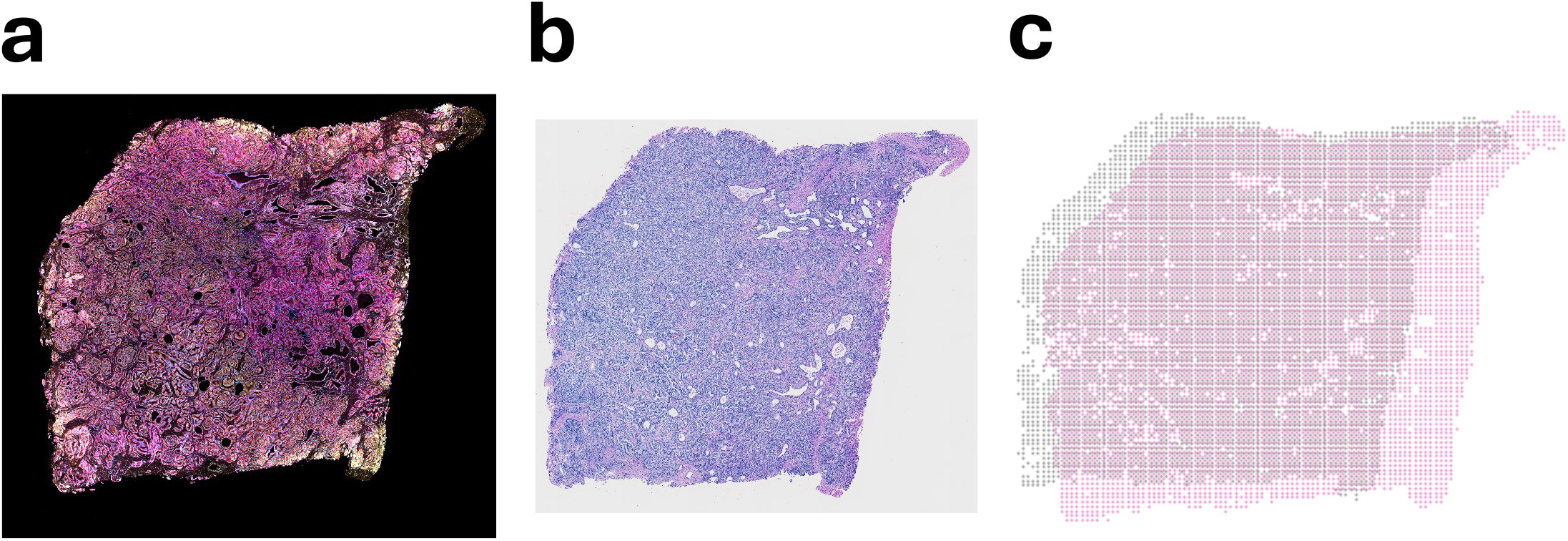
The farthest pair of adjacent slices in the HT891Z1 sample. **a**, The unaligned image of slice Z100. **b**, The unaligned image of slice Z135. **c**, Stacking of the unaligned coordinates of the two slices.

**Fig. S4:**
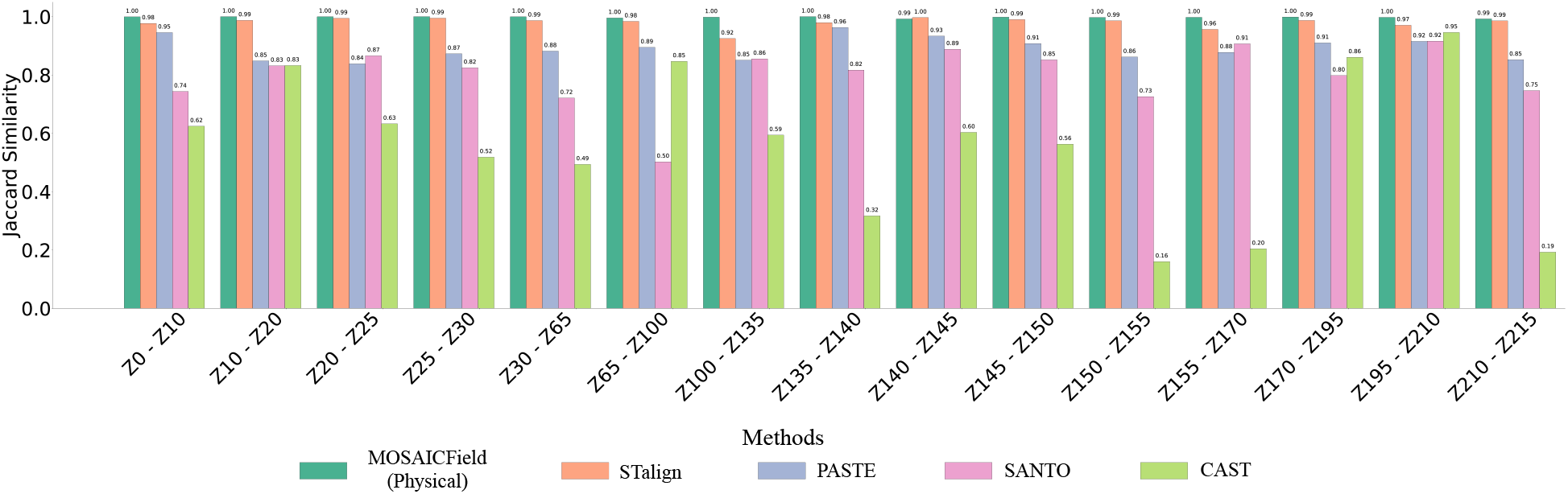
Jaccard Similarity between each pair of adjacent slices in HT891 aligned by MOSAICField physical alignment and competing methods. The alignments are applied on a global scale, aligning whole slices.

**Fig. S5:**
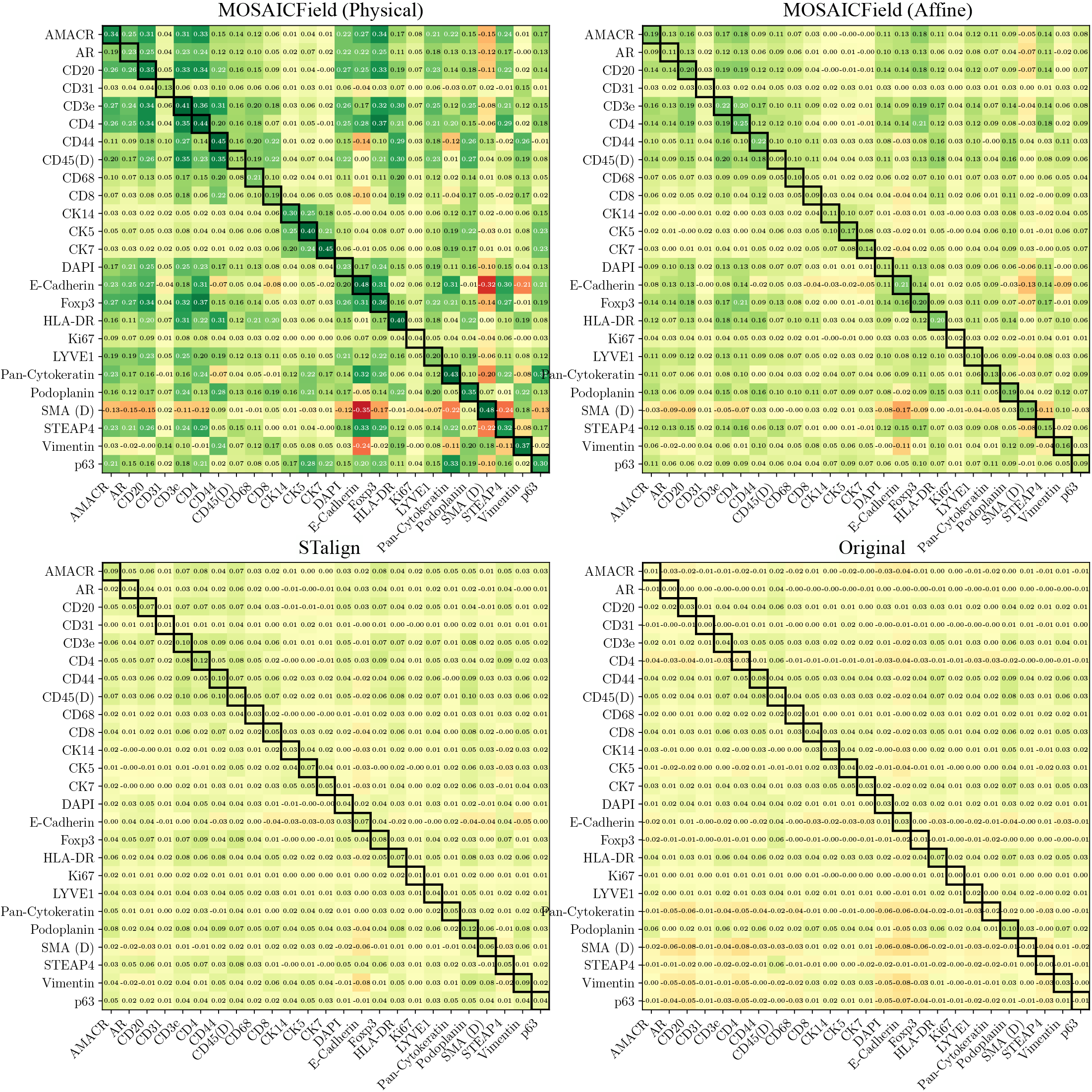
Correlations for every pair of 25 proteins in the CODEX panel between Z195 and Z210. The correlations are computed using MOSAICField physical alignment, STalign, MOSAICField affine alignment, and raw unaligned coordinates. The alignments are applied on a global scale, aligning whole slices. MOSAICField improves unimodal protein-protein correlations.

**Fig. S6:**
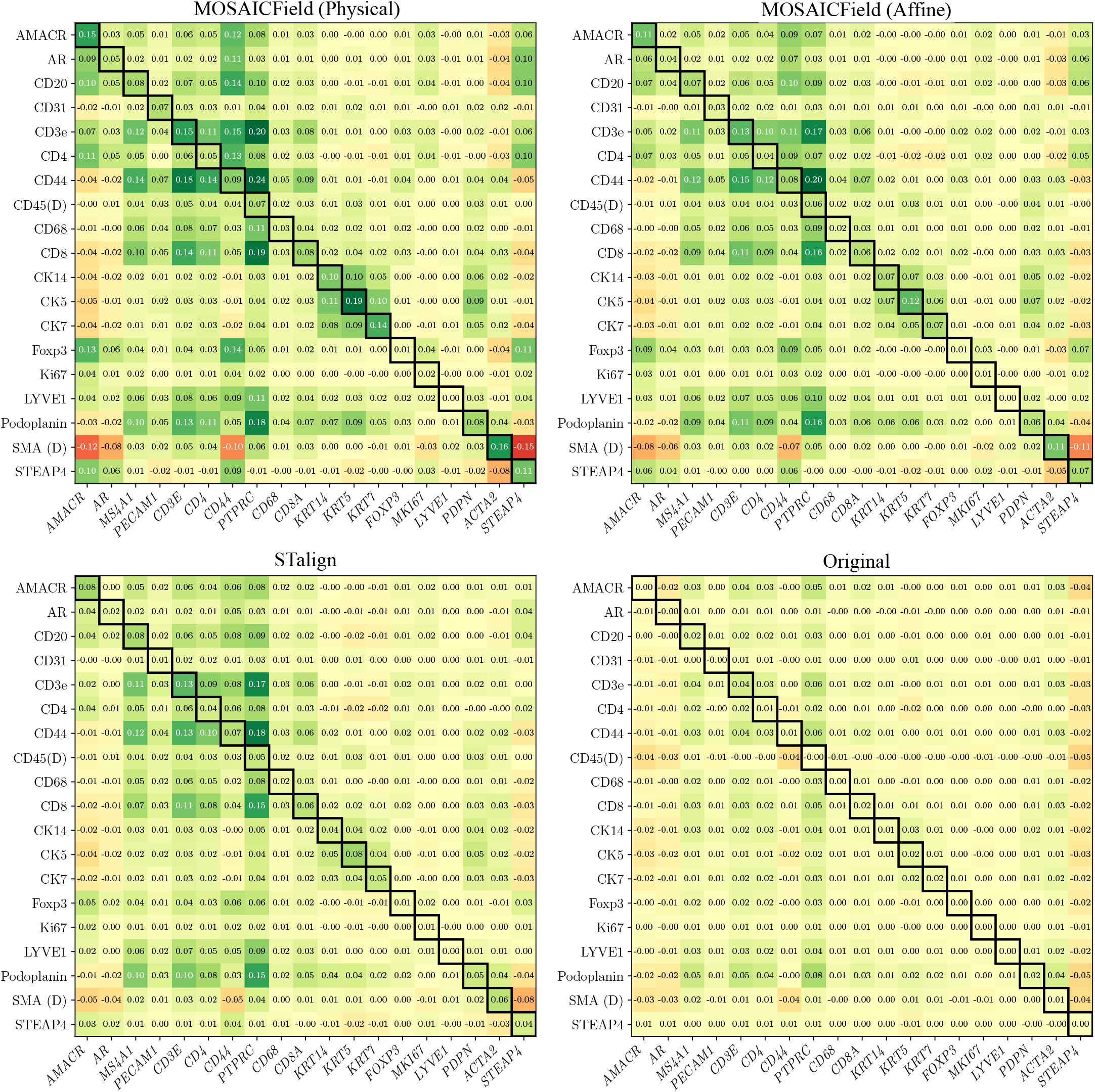
Correlations for every pair of 19 proteins in the CODEX panel and 19 coding genes in the Xenium panel between Z210 and Z215. The correlations are computed using MOSAICField physical alignment, STalign, MOSAICField affine alignment, and raw unaligned coordinates. The alignments are applied on a global scale, aligning whole slices. MOSAICField improves multimodal protein-gene correlations. Six proteins in the CODEX panel are left out because they either do not have a well-defined coding gene, or the coding gene is absent in the Xenium panel.

**Fig. S7:**
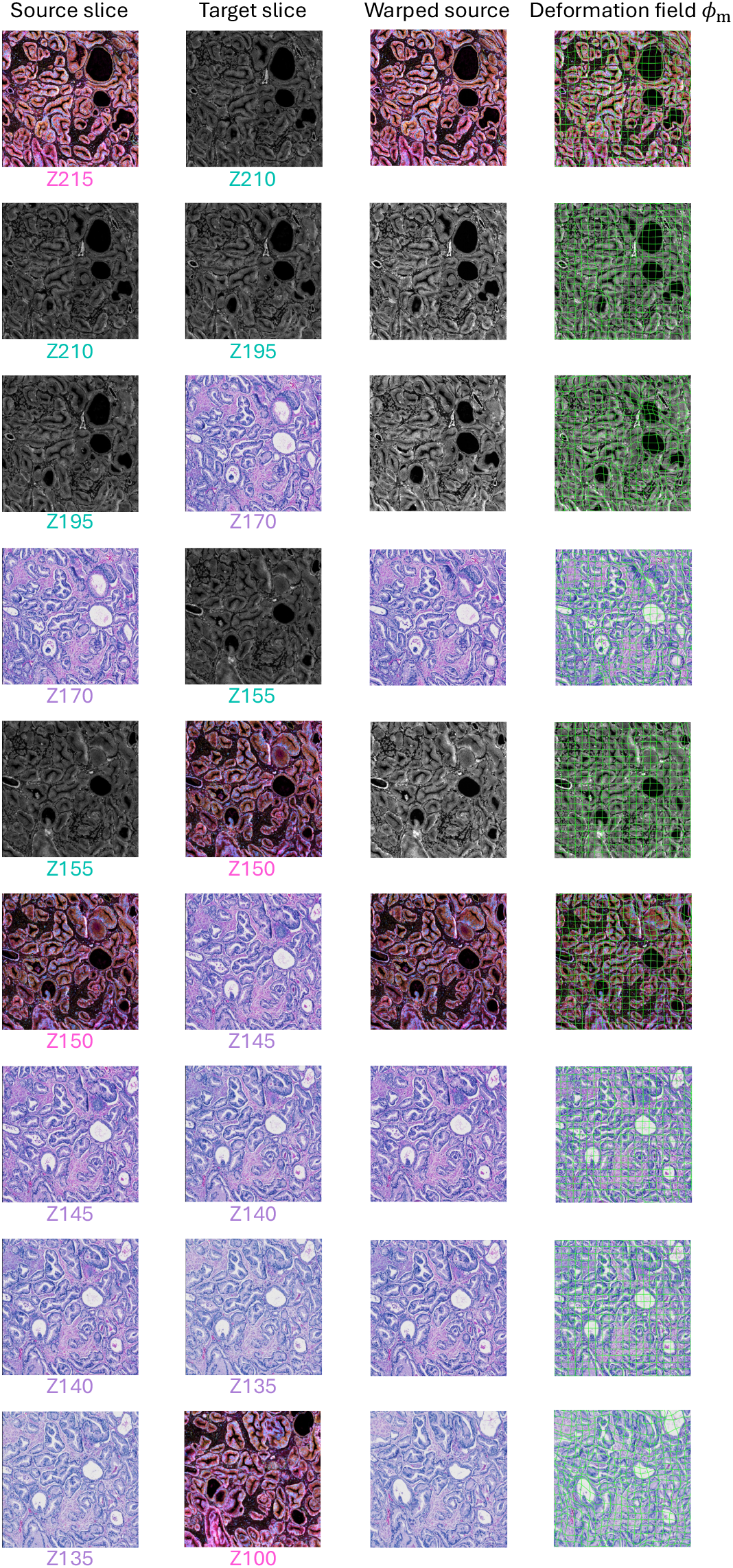
MOSAICField morphological alignment deformation field of the local region for all adjacent pair of slices in HT891 sample. From left to right column: source image, target image, warped source image and warped source image with deformation field visualized.

**Fig. S8:**
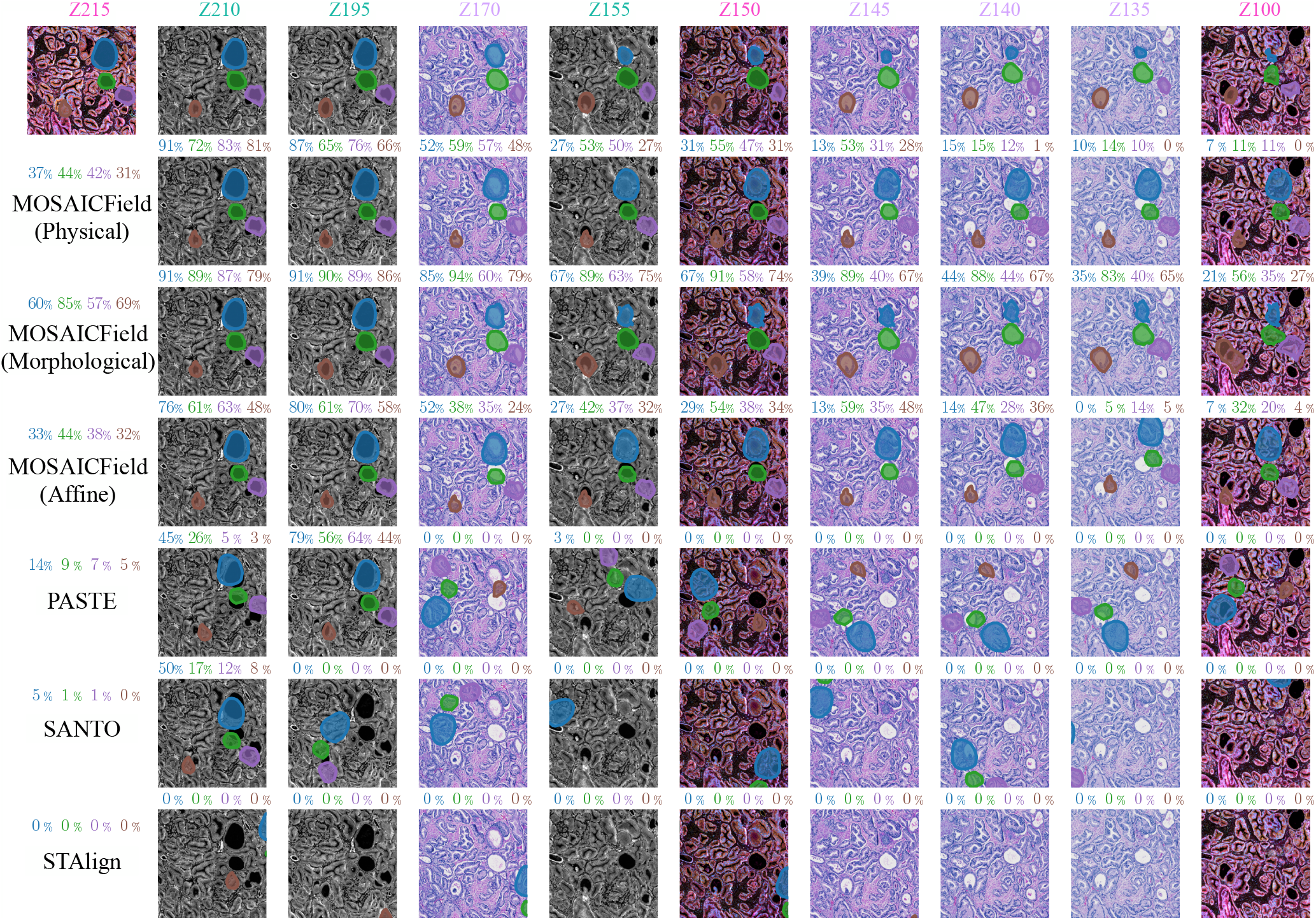
Reference manual annotation of four ductal ROIs in the local region across 10 slices and morphology tracking results of all benchmarked methods. Colored percentage values above each plot indicate Jaccard similarity of each ROI within the slice against reference annotation at the top row. Value above method names on the leftmost column indicate average Jaccard similarity across all non-reference slices.

**Fig. S9:**
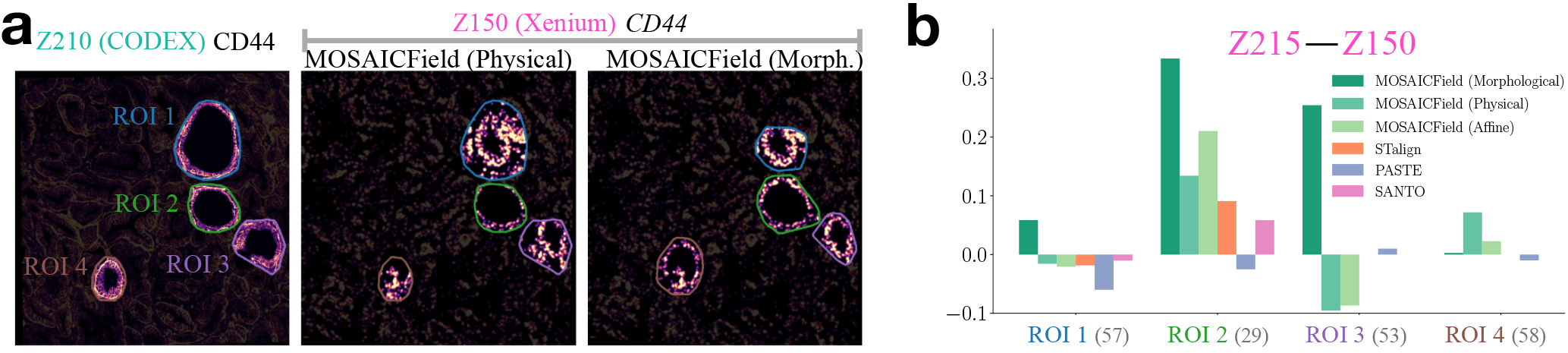
Feature holdout experiments. **(a)** We run MOSAICField morphological alignment on HT891 while holding out expression of gene *CD44* and protein CD44 when computing the alignment, then repeat the procedure as done in Figure 5A. **(b)** We run MOSAICField morphological alignment on HT891 while holding out 90% of genes in Xenium datasets, and compute average PCC over holdout genes as done in Figure 5D. In both scenarios, we observe similar results as in Figure 5 where features are not held out.

**Fig. S10:**
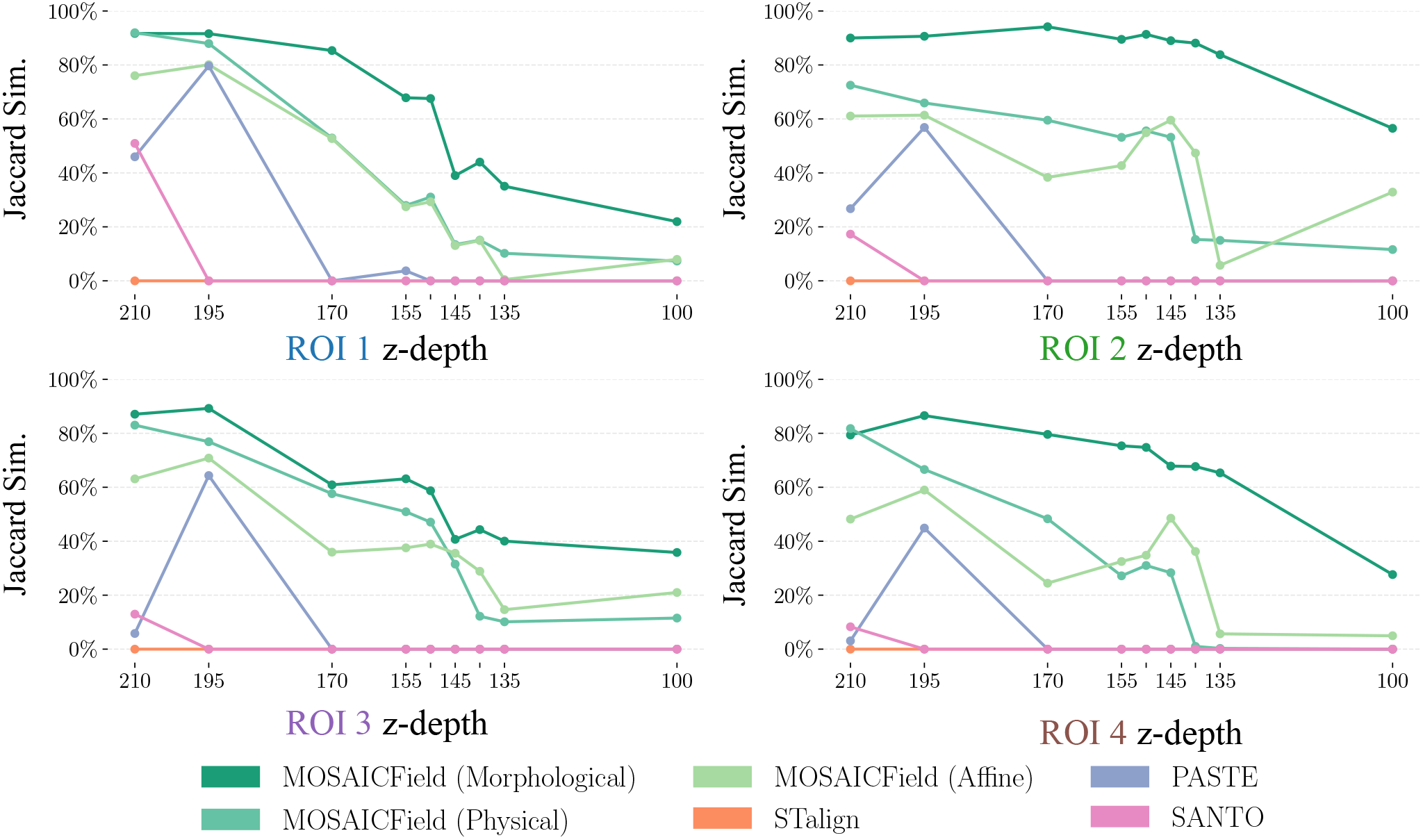
Trendline of Jaccard similarity against reference annotation for benchmarked methods. One plot for each ROI, ordered by decreasing Z (in the same way alignments are run).

**Fig. S11:**
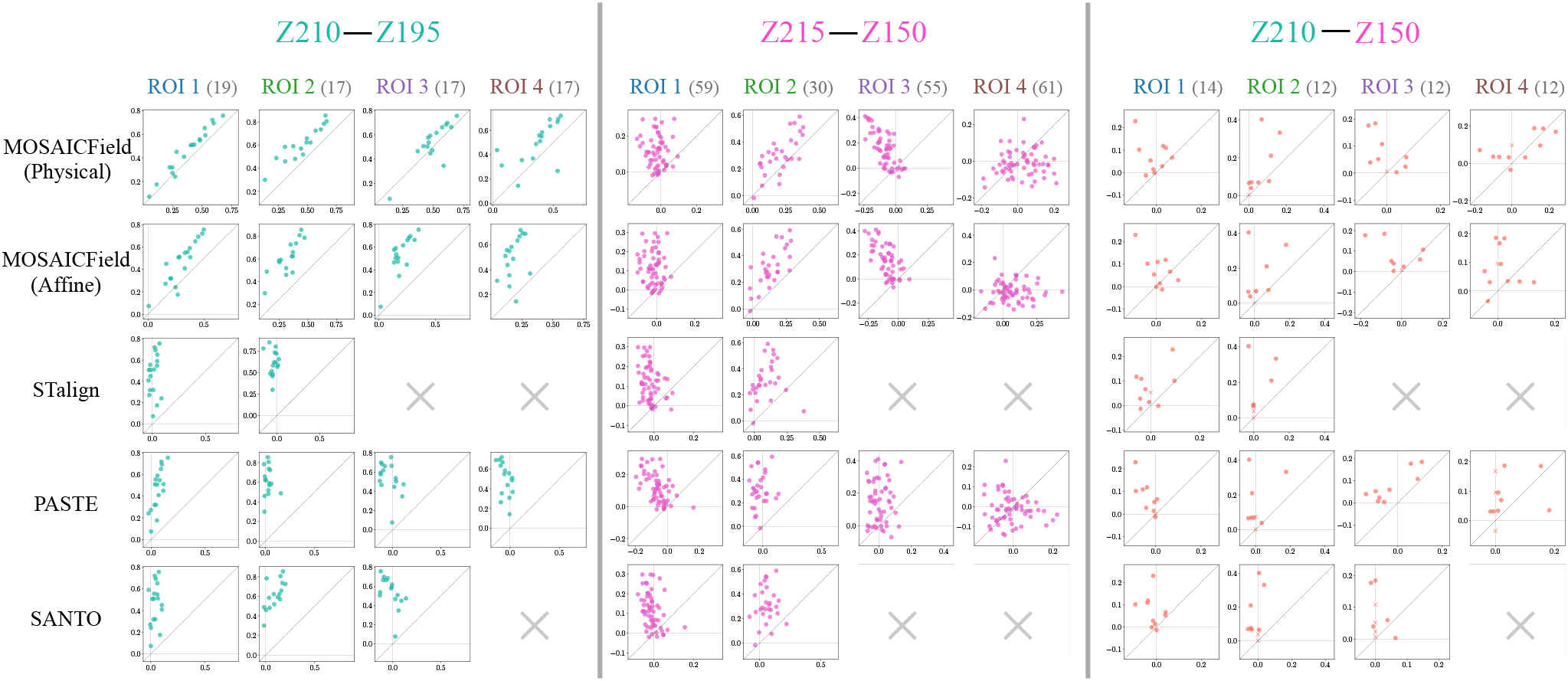
For each of the three profiled pairs and for each ROI (column), we plot the pixel-level Pearson Correlation Coefficient for MOSAICField morphological alignment (Y-axis in each subplot) against every other approach (X-axis in each subplot). Grey X in place of a subplot indicates the benchmarked method transforms this ROI fully outside the 1mm 1mm region, in which case the correlation scores represent pure noise. In case the transformed ROI has zero readout for a gene or protein, the correlation is assumed to be 0.

